# Structural Plausibility Without Binding Specificity: Limits of AI-Based Antibody-Antigen Structure Prediction Confidence Scores

**DOI:** 10.64898/2026.03.02.709004

**Authors:** Eva Smorodina, Montader Ali, Klara Kropivšek Brumat, Leonardo Salicari, Samo Miklavc, Aibek Kappassov, Chengcheng Fu, Pietro Sormanni, Ario de Marco, Victor Greiff

## Abstract

Antibody-antigen binding prediction remains a central challenge for AI-driven therapeutic discovery, particularly in discriminating cognate interactions from structurally plausible but incorrect pairings. We present a controlled, AI-method- and antibody-format-agnostic evaluation framework that measures binding specificity under realistic conditions. Using 106 experimentally determined single-chain antibody (nanobody)-antigen complexes and 11,342 shuffled non-cognate pairings, we benchmarked publicly-available state-of-the-art structure prediction methods (AlphaFold3, Boltz-2, Chai-1). Although the methods tested often generated geometrically plausible complexes, internal confidence metrics (ipTM) frequently failed to discriminate correct from incorrect pairings. Increased sampling improved structural refinement but not pairing discrimination, indicating that computational resources are better allocated across independent seeds and explicit negative controls. We conclude that internal confidence scores are not inherently calibrated to binding specificity and require validation against realistic decoys. To enable community benchmarking and method development, we release ∼1.8 million AI-generated complex structures and guidance for the benchmarks ahead.

## Introduction

Antibodies are key immunotherapeutic biomolecules characterized by antigen-specific binding. This specificity enables antibodies to identify and bind molecular targets such as tumor- or pathogen-associated antigens ^1^, underpinning a wide range of therapeutic applications and positioning antibodies as the largest class of biotherapeutics ^2^. With the continued growth of monoclonal antibody (mAb) therapeutics, there is increasing interest in developing in silico antibody discovery and design methods ^3–7^. While experimental discovery pipelines still rely heavily on antibody libraries and screening ^8^, improved prediction of paratope-epitope interactions may enable in silico discovery strategies ^3^.

Antigen recognition is primarily mediated by the antibody’s six hypervariable complementarity-determining region (CDR) loops spread over heavy and light chains that together form the paratope ^9^. Other classes of antibodies (such as nanobodies also known as VHHs^10^) have only 3 CDRs (the heavy chain loops only). These loops engage antigen surface regions (the epitope) through a combination of shape complementarity, physicochemical interactions, and conformational adaptability ^11–14^. Although recent advances in deep learning have revolutionized protein structure prediction for many molecular systems ^15–17^, accurately predicting antibody/nanobody-antigen complexes in general and identifying the correct paratope-epitope interface in particular remains challenging ^18,19^. Previous benchmarking studies report success rates of approximately 20% for antibody-antigen docking using AlphaFold-Multimer (v2.3.0) and Rosetta-based protocols ^20–23^. More recent evaluations indicate improved performance ^24,25^, with ∼35% success using a single stochastic seed and up to ∼60% success when extensive sampling (up to 1000 seeds) is combined with confidence-based ranking ^15,26^, albeit at substantial computational cost.

Recently, several state-of-the-art molecular structure prediction models have emerged, including AlphaFold2/3 ^15,27^, Boltz-1/2 ^16,28^, and Chai-1/2 ^29,30^. Several performance evaluations on diverse benchmarks spanning protein monomers, multimers, and small-molecule interactions have been made ^31–34^ and suggested that some of these models outperform earlier generations such as, for example, AlphaFold-Multimer ^35^. However, systematic evaluation of their ability to identify correct antibody-antigen interfaces and their inability to identify incorrect interactomes remains absent. Evaluating these computational complex-prediction models can highlight key metrics that can be used to discriminate cognate binding partners (“real” complexes) from non-cognate ones (“shuffled” complexes, negative controls) ^36,37^. This distinction is particularly important in discovery and screening contexts, where large numbers of candidate antibody-antigen pairings must be evaluated to separate binders from non-binders and wrong-epitope binders, with false positives proving to be a persistent challenge ^26,38,39^.

To explicitly address the problem of binding partner discrimination, we introduce a benchmarking framework that enables ground truth distinction between real and shuffled nanobody (here: VHH) and antigen complexes (Figure 1). Real complexes are defined as cognate VHH-antigen pairs extracted from experimentally solved structures, where the VHH and antigen are known biological binding partners within the same Protein Data Bank (PDB) entry. In contrast, shuffled non-cognate complexes are artificially generated, non-cognate VHH-antigen pairs created by pairing a VHH sequence from one experimental complex with an antigen sequence from a different experimental complex. These non-cognate shuffled pairs do not correspond to known biological interactions and serve as ground-truth decoys. This benchmark design allows direct comparison of computational predictions across cognate and non-cognate pairings under identical modeling conditions.

**Figure 1.**
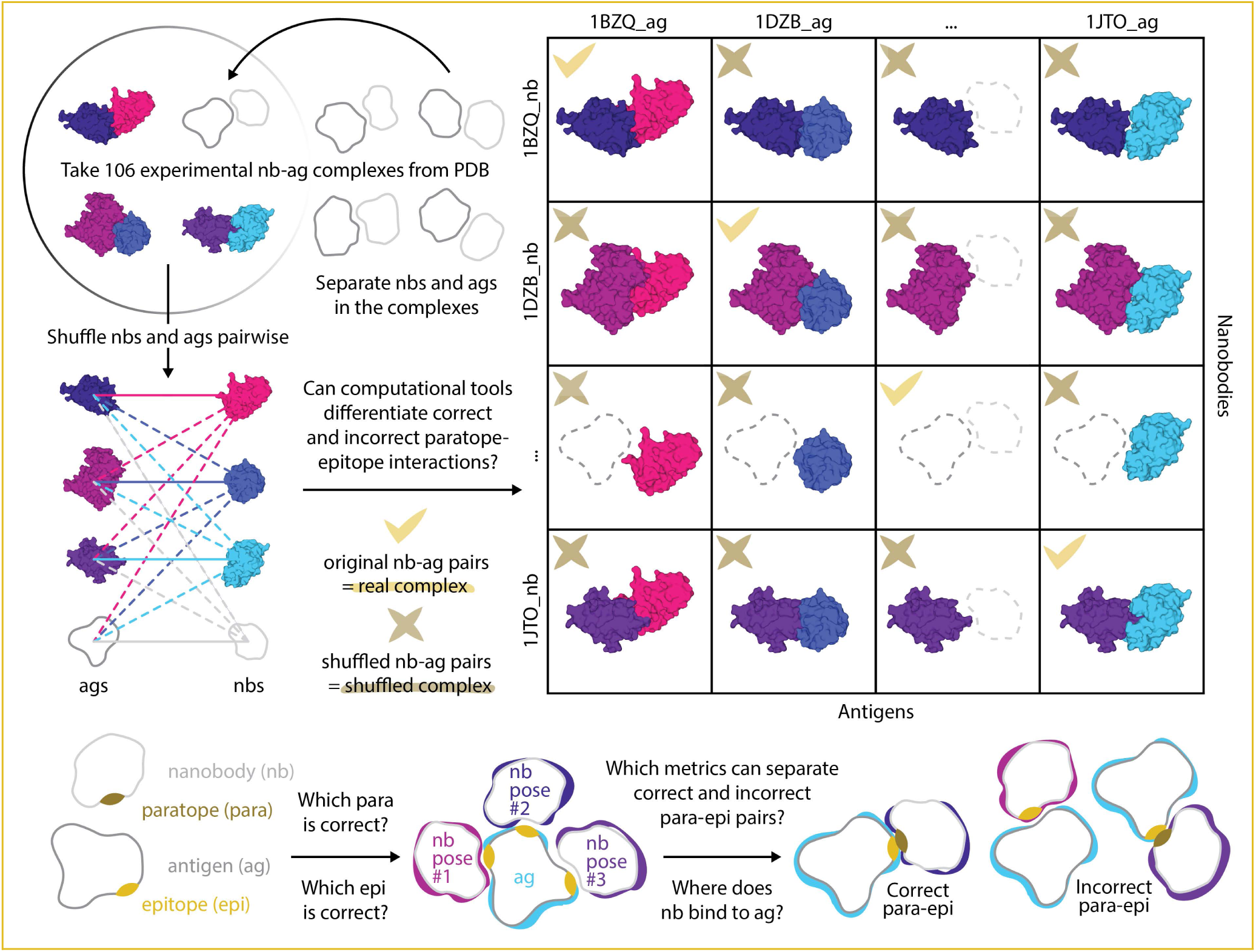
Paratope-epitope identification challenge. Schematic overview of the real-versus-shuffled nanobody-antigen pairing strategy used to assess whether computational methods can correctly prioritize cognate paratope-epitope interactions. Starting from 106 experimentally resolved nanobody-antigen complexes deposited in the PDB (left), nanobodies (VHH) and antigens are separated and then recombined pairwise. Original nanobody-antigen pairings correspond to real complexes (Supplementary Table 1), while shuffled nanobody-antigen combinations form shuffled (non-cognate) complexes. The central grid illustrates representative examples of real (✔) and shuffled (✘) pairings, highlighting cases where a nanobody paratope (“para”) is presented with the correct or incorrect antigen epitope (“epi”). Nanobodies are shown in purple/magenta, antigens in blue/cyan, with paratopes and epitopes highlighted. The benchmark tests whether computational metrics can (i) distinguish real from shuffled complexes, (ii) identify the correct epitope for a given paratope, and (iii) penalize incorrect binding modes that are geometrically plausible but biologically incorrect.

We benchmarked AlphaFold3, Boltz-2, and Chai-1 for predicting VHH-antigen paratope-epitope interactions, quantifying their ability to distinguish real cognate from shuffled non-cognate pairs. This allowed us to assess structural and interface accuracy (DockQ ^40,41^), test whether confidence metrics (e.g., ipTM ^35,42^) can discriminate real from shuffled complexes, and identify sequence/structure features linked to high or low confidence independent of pairing correctness. Overall, we present a ground-truth based benchmark pipeline that exposes current model limitations and highlights opportunities to improve antibody/nanobody-antigen complex prediction for practical screening workflows.

## Results

### Computational predictors largely fail to discriminate real from “shuffled” nanobody-antigen complexes

To assess whether publicly available state-of-the-art structure prediction methods can distinguish biologically observed nanobody-antigen interactions from shuffled complexes, we evaluated three AI-based complex structure prediction tools (AF3, Boltz-2, and Chai-1) across a panel of 561,800 (106^2^×50, 561,800×3=1,685,400 for all 3 tools) VHH-antigen pairings (Figure 2): 106 real systems (complexes) from 91 unique PDB ids and 11,130 (106^2^=11,236, 11,236-106=11,130) shuffled complexes of 50 replicates each, where a replicate is defined as a prediction sample generated by a model. The selection of 106 real VHH-antigen complexes was based on their sequence and structural diversity, size, and uniqueness (Supplementary Figure 1, Methods, Supplementary Table 1). The larger number of systems (106) compared to PDB IDs (91) is due to 15 PDB entries containing more than one VHH bound to the same antigen at different epitopes. For all complexes, information on whether they were present in the training set of the respective AI tool was also available (see below). In the following, we assume that VHHs and antigens of “shuffled complexes” are non-binders.

**Figure 2.**
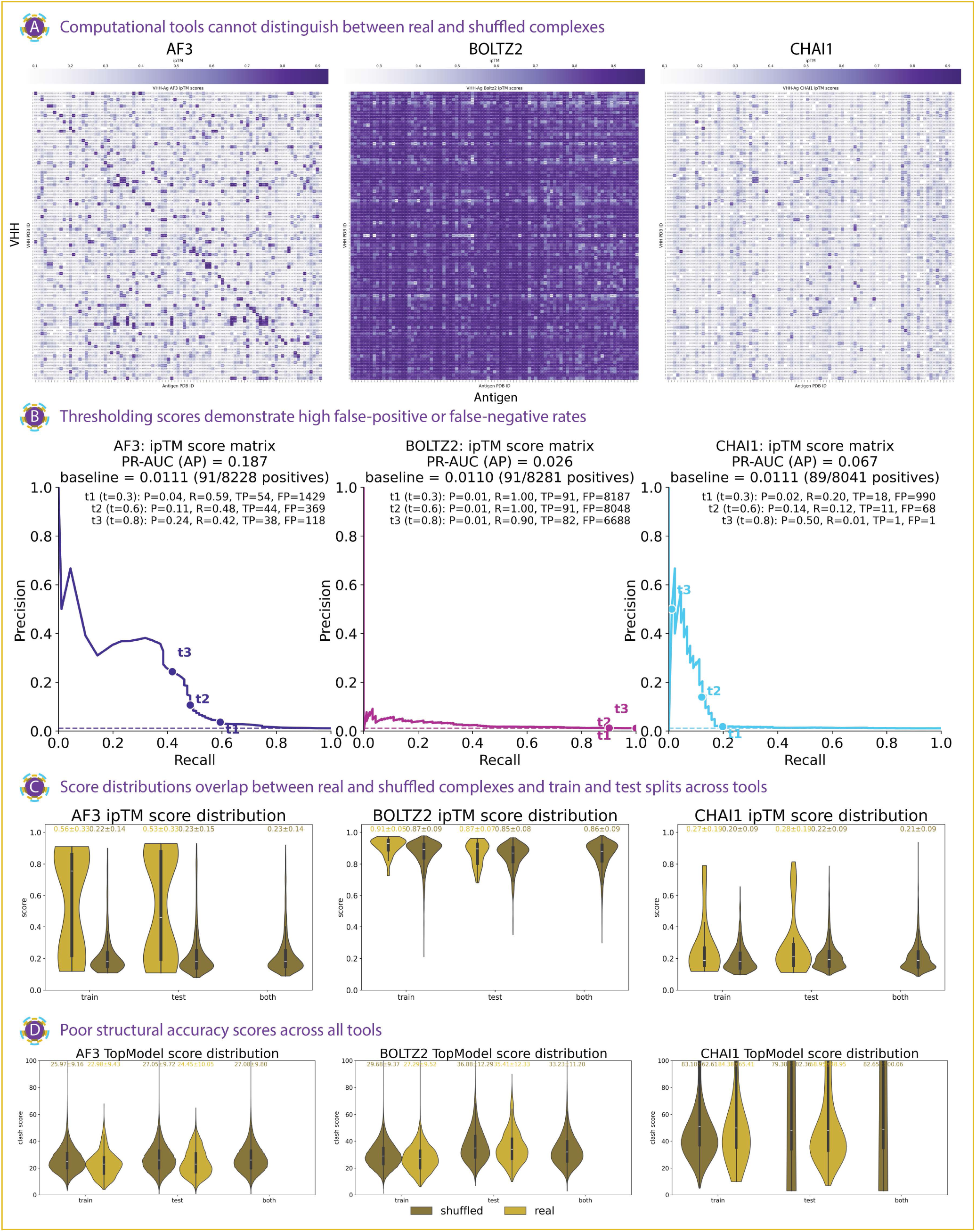
Current structure prediction tools and scores do not capture nanobody-antigen specificity. A. *Global interaction score landscapes reveal limited specificity across prediction tools.* Heatmaps show the best predicted interface confidence scores (ipTM) out of 50 replicas (structure prediction samples) per each system for all pairwise VHH-antigen combinations generated by AF3, Boltz-2, and Chai-1 (fixed seed, MSAs were generated automatically for everything except Chai-1, instead embeddings were used with Chai-1). Rows correspond to VHHs and columns to antigens. For all three tools, high-scoring interactions are broadly distributed throughout the matrices, with no clear enrichment along the diagonal corresponding to cognate (real) complexes. AF3 and Chai-1 display sparse, heterogeneous high-score patterns, whereas Boltz-2 assigns uniformly high scores across most combinations, irrespective of biological relevance. Color scales indicate ipTM values for each tool (darker color - higher values). B. *Precision-recall analysis shows limited discrimination between real and shuffled complexes.* Precision-recall (PR) curves summarize performance across all possible ipTM thresholds for AF3, Boltz-2, and Chai-1. Because positives are rare in the all-vs-all evaluation, the dashed horizontal line indicates the random baseline corresponding to class prevalence (∼0.011 precision). PR-AUC (Average Precision, AP) values quantify overall discrimination ability, with AF3 achieving the highest performance (AP=0.187), followed by Chai-1 (AP=0.067) and Boltz-2 (AP=0.026). Reference thresholds (t1-t3; ipTM ≥0.3, ≥0.6, ≥0.8) are marked on the curves to illustrate the trade-off between precision and recall: lower thresholds increase recall but introduce many false positives, whereas higher thresholds improve precision at the cost of sharply reduced recall. Across all tools, curves remain close to the random baseline, indicating limited ability to distinguish real from shuffled complexes. C. *Score distributions overlap between real and shuffled complexes across dataset splits.* Violin plots summarize ipTM score distributions for real and shuffled complexes across training, test, and mixed (both) splits, defined based on the original train-test split of each tool. Real and shuffled distributions strongly overlap for all predictors, with comparable means and variances across splits. AF3 exhibits bimodal distributions for real complexes but substantial overlap with shuffled scores, Boltz-2 assigns consistently high scores to both classes, and Chai-1 yields uniformly low scores, indicating limited discriminatory power across all methods. D. *Structural accuracy scores are non-discriminative across prediction tools.* Violin plots show the distribution of clash scores for the top-ranked predicted nanobody-antigen complex generated by AF3, Boltz-2, and Chai-1, evaluated across training-leakage, test, and mixed (“both”) dataset splits. “Both” data split refer to either the antigen or the VHH present in the training of the model. Clash scores quantify structural quality by accounting for both the number of steric clashes and protein length, with lower values indicating better geometric plausibility. For all tools, clash score distributions are broad and strongly overlapping between real and shuffled complexes, indicating that basic steric quality does not distinguish cognate from non-cognate pairings.

First, we found that interface score landscapes (ipTM) largely lack specificity for real complexes (Figure 2A). We intentionally used only 91 unique PDB entries (not 106 systems that include the same antigens with different VHHs) to avoid mixing different analytical levels and to maintain dataset simplicity. This ensured a clean comparison of whether current structure prediction tools can distinguish real from shuffled interactions, without overrepresenting antigen contexts containing multiple VHHs. All-VHH-vs-all-Ags heatmaps of predicted interface confidence scores revealed broadly similar interaction landscapes for real and shuffled complexes across all three tools. AF3 and Chai-1 produced sparse, heterogeneous score patterns with isolated high-scoring interactions distributed throughout the matrix, while Boltz-2 assigned uniformly high scores across most VHH-antigen combinations. While AF3 showed some enrichment of scores along the diagonal of the heatmap (real cognate pairs), none of the methods displayed systematic diagonal enrichment compared to the shuffled complexes, indicating, qualitatively, a lack of specificity for biologically correct pairings.

To quantify whether cognate (diagonal) pairs are enriched relative to shuffled (off-diagonal) pairs (Figure 2A), we evaluated each tool using precision-recall (PR) curves and their area under the PR curve (PR-AUC; Average Precision, AP), which are less sensitive to class imbalance than ROC-based metrics and avoid reliance on an arbitrarily chosen cutoff (Figure 2B). Across all three predictors, PR performance remained low. AF3 achieved the highest PR-AUC (AP = 0.187), followed by Chai-1 (AP = 0.067) and Boltz-2 (AP = 0.026). Because positives are rare in this all-vs-all setting, the random baseline in PR space equals the positive prevalence and is correspondingly low (baseline precision ≈ 0.011; ∼89-91 positives out of ∼8041-8281 evaluated pairs, depending on the tool after excluding missing values). Thus, while AF3’s AP is above baseline, all tools remain far from reliably separating real from shuffled complexes, consistent with the weak diagonal enrichment observed in the heatmaps.

For interpretability, we additionally overlaid three reference thresholds ^42,43^ (ipTM≥0.3, 0.6, 0.8, denoted t1-t3, chosen according to the scoring metric literature ^42,43^) onto the PR curves (Figure 2B). These markers illustrate the expected threshold trade-off: lower thresholds increase recall at the expense of precision (many false positives), whereas higher thresholds improve precision but sharply reduce recall (many false negatives). This effect is especially pronounced under extreme class imbalance, making single-threshold summaries (e.g., confusion matrices, F1, specificity) highly threshold-dependent and potentially misleading when comparing tools. In Boltz-2, uniformly elevated scores drive near-maximal recall at low-to-intermediate thresholds but with extremely low precision, reflecting widespread false positives rather than meaningful discrimination. Overall, PR curves and PR-AUC confirm that none of the evaluated tools achieve robust separation of real nanobody-antigen complexes from shuffled mismatches.

Next, we asked to what extent scores differed between pairs that belong to the train or test sets within each dataset split (Figure 2C). Dataset splits were defined using the original train-test splits of each tool: complexes were labeled as “train” when both the VHH and antigen were in complex (real) and present in the training set of that model respectively (as of its training set date-cutoff); as “test” when both were in complex (real) but not seen by the model, as “both” when one of the two binding pairs was drawn from the training set and the other from the test set (shuffled). For this analysis, we used the following datasplits: Chai-1- 25 train and 81 test systems; Boltz-2 - 64 train and 42 test systems; AF3 - 30 train and 76 test systems. Violin plots summarizing ipTM score distributions confirm substantial similarity of interface scores between real and shuffled complexes for all tools and across training, test, and mixed (both) splits. AF3 showed bimodal distributions for real complexes, with a subset of high-scoring interactions, but shuffled complexes still occupy overlapping score ranges. Boltz-2 assigns consistently high ipTM values to both real and shuffled interactions (mean ∼0.85-0.91), with minimal separation. Chai-1 yielded uniformly low scores for both classes, again with nearly identical distributions. Across all tools, mean scores and variances were comparable between real and shuffled complexes, indicating that score magnitude alone does not encode interaction authenticity.

We used TopModel’s clash score ^44^ to evaluate the structural quality across real and shuffled nanobody-antigen complexes (Figure 2D). The clash score accounts for both the number of steric clashes between all atoms and protein length, with lower values indicating better geometric quality. Across all tools and dataset splits, clash score distributions were broad and overlapped between real and shuffled complexes. For AF3, top-ranked (highest ipTM) models show mean clash scores of 25.7±9.1 (train), 26.7±9.8 (test), and 26.8±9.7 (both) for shuffled complexes, compared with 22.7±9.4, 24.2±9.9, and 26.8±9.7 for real complexes across the same splits. Boltz-2 produced systematically higher clash scores than AF3, with shuffled complexes averaging 30.0±9.5 (train), 36.1±12.4 (test), and 33.0±11.3 (both), while real complexes average 27.9±10.7, 34.8±12.2, and 32.9±11.3, respectively. Chai-1 exhibits markedly poorer structural quality by this metric. Mean clash scores for Chai-1 were 4-5 times higher and more variable, reaching ∼80-90 on average, with extreme dispersion (e.g., 62.0±65.0 and 90.7±125.2 in the test split). These elevated and highly variable scores were observed for both real and shuffled complexes, indicating frequent steric clashes in top-ranked predictions and a lack of effective internal filtering for geometric plausibility. Importantly, real complexes did not systematically achieve lower clash scores than shuffled complexes in any tool or dataset split. In several cases, shuffled complexes displayed comparable or even slightly lower average clash scores than real complexes, despite being biologically incorrect.

To assess whether different prediction tools agree on which nanobody-antigen pairs are confident or uncertain, we directly compared ipTM score matrices generated by AF3, Boltz-2, and Chai-1 (Figure 2A). Global correlations between flattened score matrices were uniformly low (pearson r=0.14 for Chai-1-Boltz-2, r=0.18 for Chai-1-AF3, and r=0.13 for Boltz-2-AF3), indicating low agreement across tools in their assessment of interaction confidence.

Per-system cross-tool correlation analysis, which examines whether tools predict the same systems with similar levels of confidence or lack thereof, further revealed substantial heterogeneity across both VHHs and antigens. For each pair of tools, many systems exhibited weak or even negative correlations, highlighting disagreement in how individual VHHs or antigens are scored (Supplementary Figure 2A). Importantly, these discrepancies were not confined to a small subset of problematic systems but were broadly distributed across the benchmark, suggesting that tool-specific inductive biases strongly shape confidence assignment. Outlier analysis reinforces this conclusion. High-confidence “shuffled” outliers identified by one tool rarely overlapped with those from another. Specifically, only four shuffled outliers overlapped between Chai-1 and Boltz-2, fourteen between Chai-1 and AF3, and just one between Boltz-2 and AF3, despite all tools being evaluated on the same set of VHH-antigen pairings. Representative examples illustrate the magnitude of these discrepancies. For instance, the system (4NC2, 7R24) was assigned high confidence by Boltz-2 (ipTM≈0.81) but substantially lower confidence by Chai-1 (ipTM≈0.60), while (9EMY, 8UKV) achieved high confidence in Chai-1 (ipTM≈0.73) but was scored markedly lower by Boltz-2 (ipTM≈0.44). Similarly, several systems such as (7TGF, 6OBO) and (8K4Q, 8EW6) were consistently high-confidence outliers in AF3 (ipTM≥0.8) yet did not emerge as outliers in the other tools (Supplementary Table 2).

Together, these results show that both ipTM-confidence and structural accuracy remain limited across tools, with AF3 producing the lowest average clash scores, Boltz-2 intermediate values, and Chai-1 the poorest overall structural quality. However, even the best-performing method by this metric fails to reliably discriminate real nanobody-antigen complexes from shuffled mismatches. This inability to distinguish real from shuffled complexes persists across training, test, and combined datasets, indicating that the observed overlap is unlikely to result from overfitting or data leakage. Instead, it reflects a limitation of current structure-based confidence scores and simple structural quality measures, which capture generic interface properties but ultimately fail to reflect biological correctness in nanobody-antigen complex prediction.

### Cross-tool score agreement is weak and sampling improves structure but not confidence alignment

Next we focused on a thorough evaluation of the real complexes (the diagonal systems of the Figure 2A matrices) against their experimental references. We asked whether models “know” when they fail - that is, whether confidence scores (ipTM) reliably indicate structural prediction quality (DockQ) under different sampling regimes (Figure 3). Confidence calibration - the correspondence between predicted confidence and actual quality ^45,46^ is critical for practical screening workflows where ground-truth access to structural information is absent. Thus, confidence calibration analysis was only performed on the real complexes, as shuffled pairings lack experimental reference structures against which to compute DockQ ^41^. We found that confidence calibration differs systematically across tools when we display correlation of of DockQ versus ipTM for the best, initial (sample₀), and worst of 50 predictions (Figure 3A).

**Figure 3.**
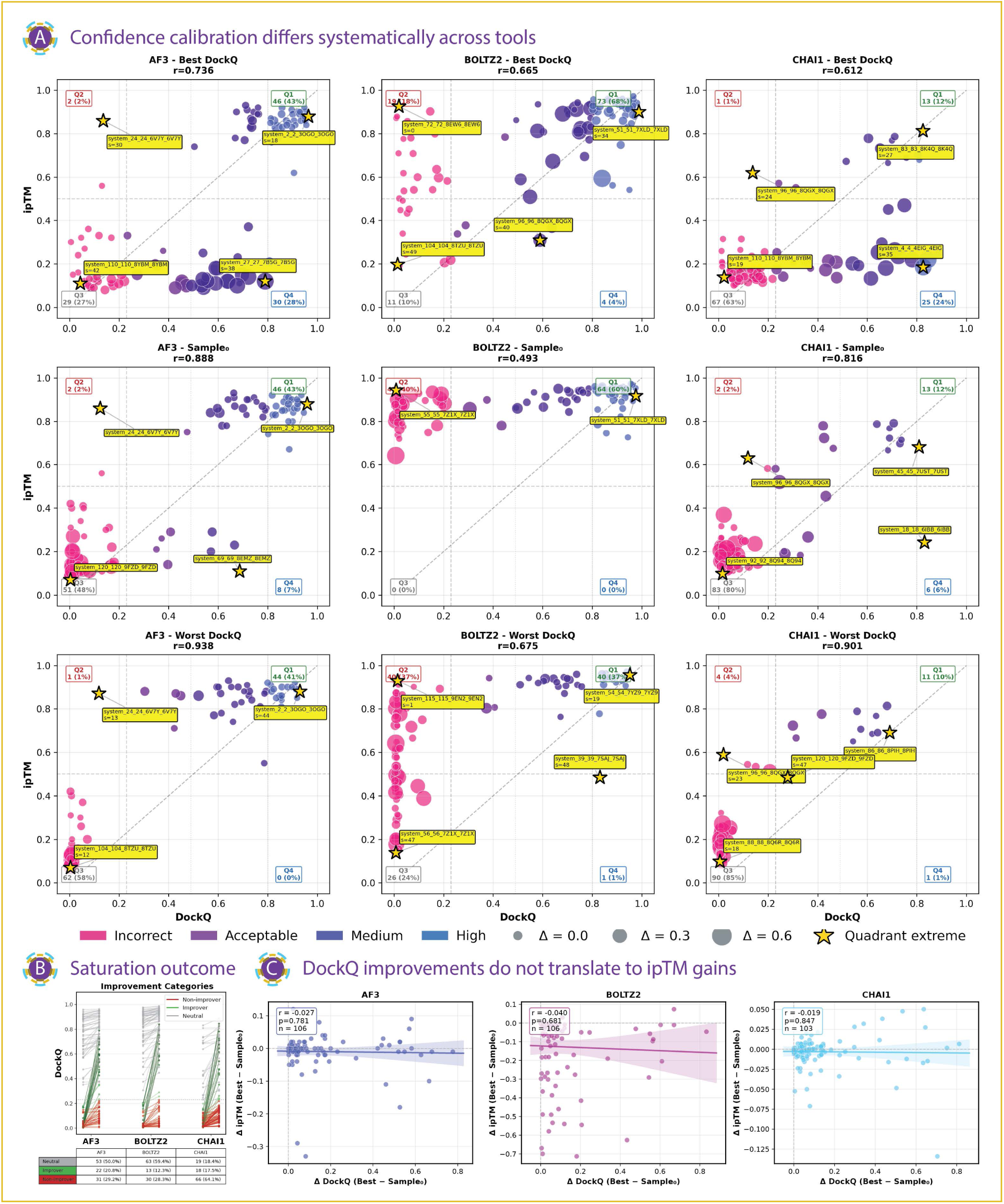
Cross-tool disagreement and confidence calibration of nanobody-antigen docking predictions. A. *Confidence calibration differs systematically across ML structure prediction tools.* Scatter plots show DockQ versus ipTM for AF3, Boltz-2, and Chai-1 under three sampling regimes: best DockQ, initial sample (sample₀), and worst DockQ. Points are colored by the official DockQ quality category (incorrect, acceptable, medium, high) and sized according to the saturation range (ΔDockQ from sample₀ to best). Yellow stars mark quadrant extremes, highlighting overconfident failures (high ipTM, low DockQ) and underconfident successes (low ipTM, high DockQ). Dashed lines indicate DockQ=0.23 (acceptable quality threshold) and ipTM=0.5 (confidence threshold). pearson correlation coefficients (r) are shown for each panel. AF3 exhibits the strongest calibration, Boltz-2 shows systematic overconfidence, and Chai-1 displays systematic underconfidence, with calibration degrading substantially in low-sampling regimes. B. *Saturation outcome categories reveal when sampling rescues docking quality.* Line plots show per-system trajectories from the initial prediction (sample₀) to the best of the sampled predictions, grouped into three outcome classes: non-improvers (red) remain below the acceptable-quality threshold (DockQ<0.23) across all samples; improvers (green) are rescued from incorrect to acceptable quality (cross DockQ=0.23); and neutral systems (grey) are already acceptable at sample₀ (DockQ≥0.23) and may further refine with sampling. Systems with known structural artifacts and runs that failed to complete were excluded (14 systems; see Methods). A comprehensive list of improvement categories is provided in Supplementary Table 4. C. *DockQ gains from sampling do not translate into confidence-score gains.* Scatter plots show, for each system and model, the relationship between ΔDockQ (best - sample₀) and ΔipTM (best - sample₀), with linear regression fits and 95% confidence intervals. Near-zero correlations indicate that confidence scores largely fail to track structural quality improvements achieved through saturation sampling (AF3: N=106, r=-0.027; Boltz-2: N=106, r=-0.040; Chai-1: N=103, r=-0.019), consistent with confidence being “locked in” to early trajectory choices rather than reflecting refinement.

To interpret calibration patterns, we defined four diagnostic quadrants based on thresholds of DockQ=0.23 (acceptable quality) and ipTM=0.5 (confident prediction): Q1 (confident and correct), Q2 (overconfident failures), Q3 (uncertain and incorrect), and Q4 (underconfident success) (Supplementary Table 3). Among the three methods, AF3 exhibited the strongest calibration, with a pearson correlation of r=0.736 for best-DockQ structures and relatively few overconfident failures (Q2 - 2%, Q4 - 28%). AF3 maintained high correlation even at the single-sample level (r=0.888 for sample₀), indicating comparatively robust confidence assignment without extensive sampling. In contrast, Boltz-2 was systematically overconfident. Although its best-DockQ calibration remained moderate (pearson r=0.665), a substantial fraction of predictions fell into the overconfident failure regime (Q2 - 18%), where ipTM is high despite poor structural quality. This effect was exacerbated for single-sample predictions, where correlation dropped sharply (pearson r=0.493), highlighting the unreliability of Boltz-2 confidence scores in low-sampling screening workflows. Conversely, Chai-1 exhibited systematic underconfidence. Although a subset of predictions achieved high structural quality (DockQ≈0.4-0.9), these models were frequently assigned low ipTM scores, resulting in a substantial fraction of underconfident successes (Q4; 24%). Consequently, the relationship between confidence and best-case structural quality was weaker for Chai-1 (best-DockQ vs. ipTM, r=0.612), meaning that confidence-based filtering would exclude many structurally accurate predictions.

Extreme quadrant cases further illustrate observed failure modes. Overconfident failures (Q2) represented geometrically poor structures assigned high confidence (e.g., AF3 system_24_24_6V7Y), whereas underconfident successes (Q4) correspond to high-quality complexes that would be missed by thresholding (e.g., AF3 system_27_27_7B5G). These cases provide concrete examples of how confidence scores can diverge from structural reality in both directions (See Supplementary Table 3 for the classification of all systems in the four quadrants).

Next, we found that sampling improves structure but not confidence alignment. Across all three tools, saturation sampling improved best-case DockQ (Supplementary Figure 3A), confirming that additional sampling can refine geometry. DockQ improvement distributions are right-skewed for all models, with AF3 and Boltz-2 exhibiting the largest absolute gains, while Chai-1 showed smaller but consistent improvements. The proportion of systems achieving acceptable or higher quality (DockQ>0.23) increased substantially after saturation sampling (Figure 3B, Supplementary Figure 3B). For AF3, incorrect predictions decreased by nearly 20%, with Boltz-2 showing a comparable reduction in incorrect predictions and a marked increase in medium- and high-quality outcomes; in contrast, Chai-1 remains dominated by incorrect predictions despite sampling. This reflects systems “rescued” above the quality threshold through sampling alone (the “improvers”, Figure 3B). However, the relationship between structural improvement and confidence during saturation sampling remains weak. Whilst all three models showed significant improvements in DockQ scores (Wilcoxon p<0.001; Supplementary Figure 3C), paired analyses showed no significant improvement in ipTM for AF3 or Chai-1 (p = 0.179 and p = 0.222, respectively), and although Boltz-2 showed a statistically significant change (p < 0.001),the ipTM score values slightly decreased (Supplementary Figure 3D). In particular, correlations between changes in DockQ and changes in ipTM were near zero across all models (r=-0.027, -0.040, and -0.019 for AF3, Boltz-2, and Chai-1, respectively; Figure 3C), indicating that confidence scores were largely insensitive to structural improvements achieved through sampling; they remain “locked-in” to the initial structural hypothesis.

To assess whether confidence scores yielded consistent rankings across tools, we compared ipTM values for each system (Supplementary Figure 3E). Cross-model correlations were modest (pearson’s r = 0.29-0.39), and importantly, higher confidence in one model did not reliably correspond to higher structural quality, as measured by DockQ. From a screening perspective, this behavior has important implications. Single-sample predictions from Boltz-2 are frequently overconfident relative to achieved DockQ, whereas AF3 displays comparatively better calibration even prior to saturation sampling. Nonetheless, for all tools, confidence does not reliably track structural refinement, limiting its utility as a post-sampling ranking criterion.

Finally, we examined prediction consistency by measuring epitope-paratope variation across 15 antigens, each crystallized with two distinct VHHs binding distinct epitopes (Supplementary Figure 4A, Methods). This provides a natural test of whether models that succeed with one VHH can generalize to alternative binders of the same antigen. AF3 produced acceptable predictions (DockQ for both pairs >0.23) in 10 out of 15 cases (67%), compared to 6/15 (40%) for Boltz-2 and 2/15 (13%) for Chai-1 (Supplementary Figure 4B). Boltz-2 and, especially, Chai-1 more frequently succeeded with only one VHH or failed on both partners (Supplementary Figure 4A), suggesting that AF3 better captures the underlying antigen structure independently of the specific paratope geometry.

To summarize, we show that confident failures are largely tool-specific rather than reflecting a shared set of universally difficult or ambiguous systems. The low overlap in outlier systems and the weak agreement in confidence-based rankings across predictors highlight the absence of a common confidence landscape, underscoring the risk of relying on any single model’s confidence scores for candidate selection, particularly after saturation sampling, where structural quality improves, but confidence remains misaligned.

### Structural accuracy scores are uniformly poor, while interface contacts do not reflect specificity

We next evaluated whether predicted nanobody-antigen complexes recovered the experimentally defined antigen epitope and whether this information can discriminate real from shuffled pairings (Figure 4). DockQ provides a stringent, structure-level measure of docking correctness by comparing a predicted complex to the experimentally solved reference, and therefore is only defined for cognate (“real”) complexes that have a ground-truth structure. In our analysis, DockQ can be performed only on real pairs, whereas shuffled pairs lack a native reference against which to compute DockQ. However, DockQ can remain low even when a model contacts a broadly correct antigen surface region, because it penalizes global pose errors and interface misalignment. To evaluate interface recovery in a way that (i) is directly interpretable in terms of epitope footprint and (ii) can be applied uniformly across both real and shuffled pairings, we therefore quantify “epitope recall” as the fraction of experimentally defined antigen epitope residues recovered by the predictions, counting a residue as recovered only if it forms a VHH contact in at least 50% of stochastic replicate predictions (n=50) (Supplementary Figure 2B). This ensemble-consensus definition intentionally emphasizes reproducible contact preferences over single-sample noise, but it also collapses geometry to a residue set and does not penalize additional (non-epitope) contacts. High epitope recall does not necessarily imply a correct docking mode, but provides insight into what epitope regions are relatively more likely to be hit in predicted complexes. Epitope recall was quantified as the fraction of experimental epitope residues recovered by model predictions, where a residue was considered recovered if it contacted the VHH in at least 50% of stochastic replicate predictions (n=50). Across all three tools, epitope recall values were generally low and sparsely distributed, with substantial overlap between real and shuffled complexes (Figure 4A, Supplementary Figure 2B). Although real complexes lie along the diagonal of the epitope recall matrices, off-diagonal (shuffled) complexes frequently exhibited comparable levels of epitope recovery. This indicates that models often identify plausible antigen-contact regions even in non-cognate pairings, limiting the specificity of epitope-based discrimination. Consistent with this observation, direct comparison of epitope recall distributions showed that shuffled complexes frequently overlapped or exceeded the recall observed for real complexes across all models and data splits (Figure 4B). Focusing on the cognate pairings and whether the respective models recovered at least one residue from the true epitope (epitope recall >0), AF3 hit the most epitopes (at least one residue in the ground truth experimental epitope) in the test set of complexes (n=29 out of 76), followed by Chai-1 (n=26 out of 81) and Boltz-2 (n=17 out of 42).

**Figure 4.**
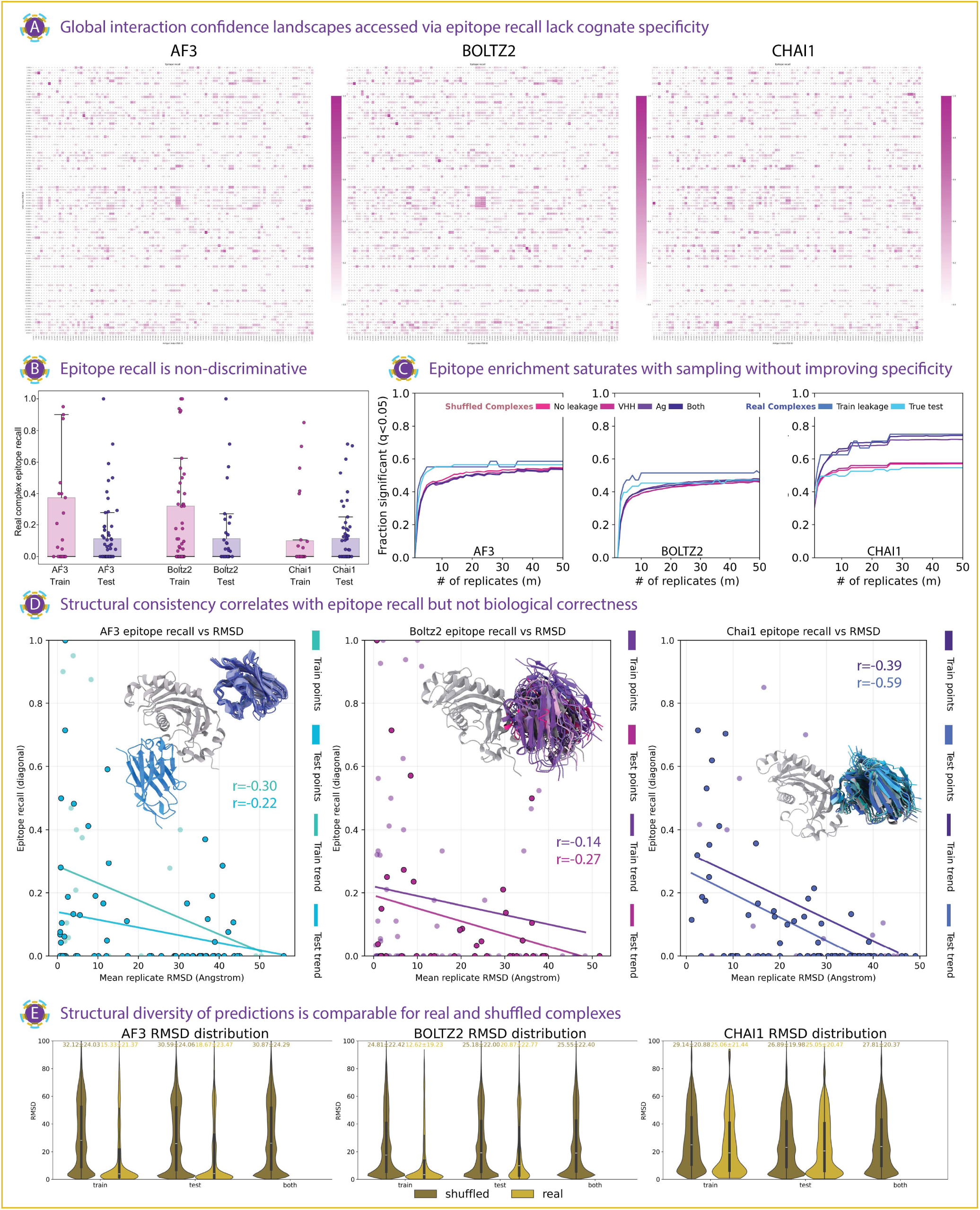
Epitope recovery and enrichment fail to distinguish cognate from non-cognate nanobody-antigen interactions. **A.** Epitope recall across real and shuffled complexes. Heatmaps show epitope recall for predicted nanobody-antigen complexes, defined as the fraction of experimentally determined epitope residues recovered by the model according to the equation shown. An epitope residue is considered recovered if it is in contact with the VHH in at least 50% of the n=50 stochastic replicate predictions. Diagonal entries correspond to real (cognate) nanobody-antigen pairs, whereas off-diagonal entries represent shuffled (non-cognate) complexes. Across all models, shuffled complexes frequently achieve epitope recall comparable to real complexes, indicating limited specificity of epitope recovery. B. *Epitope recall across real and shuffled complexes.* Distribution of epitope recall across real and shuffled complexes. Boxplots summarize epitope recall values for real and shuffled VHH-antigen pairs, stratified by train-leakage and true test sets for each model. Shuffled complexes frequently overlap or exceed the recall observed for real complexes, demonstrating that high epitope recall alone does not reliably distinguish cognate interactions. The number of points when epitope predicted correctly (points >0): AF3: train - n=12, test - n=29, Boltz-2: train - n=28, test - n=17, Chai-1: train - n=9, test - n=26. C. *Epitope enrichment significance saturates rapidly with stochastic sampling.* Line plots show the fraction of VHH-antigen pairs declared significantly enriched (FDR < 0.05) as a function of the number of stochastic predictions per complex, up to n = 50. Six distributions are evaluated per model: real complexes split into train-leakage and true test sets, and shuffled complexes split according to whether the VHH, the antigen, both, or neither were present in the training data. Across all three models, enrichment significance increases rapidly and saturates by approximately 10-15 replicates, with shuffled complexes exhibiting saturation behavior comparable to real complexes. Notably, Chai-1 shows near-complete saturation for shuffled complexes at low replicate counts, indicating a strong epitope-seeking bias that reduces the discriminative value of enrichment-based metrics. These trends suggest that increasing ensemble size primarily amplifies pre-existing contact biases rather than improving identification of true cognate interactions. D. *Structural consistency correlates with epitope recovery but not interaction correctness.* Scatter plots show the relationship between epitope recall and structural consistency across predictions, quantified as the mean pairwise RMSD of all n=50 VHH replicates after superposition on the antigen (Cα atoms only). A case study of the real complex of PDB 9ETJ is shown-predicted by the different models. The first 10 structures from the 50 replicates are shown, where the predicted complexes are superimposed on the antigen from the first predicted complex, and the predicted nanobodies are shown complexed to that antigen respectively. Points correspond to individual VHH-antigen complexes, with linear fits shown separately for train-leakage and true test sets. Reported values indicate pearson correlation coefficients. Across all models, epitope recall is negatively correlated with RMSD, indicating improved epitope recovery for complexes predicted with consistent orientations; however, this relationship holds for both real and shuffled complexes and does not imply correct paratope-epitope pairing. E. *Structural diversity of predicted complexes.* Structural diversity of predicted complexes. Violin plots show RMSD distributions for real, shuffled, and mixed (“both”) complexes across train-leakage and true test sets for each model. Similar RMSD distributions between real and shuffled conditions indicate that structural consistency alone is insufficient to discriminate cognate from non-cognate nanobody-antigen interactions.

Epitope recall scores were then correlated with ipTM scores of each model respectively (on the cognate pairings alone) to assess whether each model’s internal confidence tracks interface recovery. Across cognate pairs, ipTM was only weakly associated with epitope recall for AF3 (pearson and spearman of 0.18 and 0.15 respectively) and Boltz-2 (pearson and spearman of 0.08 and 0.14 respectively), indicating that high ipTM scores frequently occur even when the predicted interface fails to recover the experimental epitope. In contrast, Chai-1 exhibited a moderate positive relationship between ipTM and epitope recall (pearson and spearman 0.46 and 0.48 respectively), suggesting the model’s confidence signal better reflects antigen side-contact recovery than the other two models. Overall, the dispersion of the correlations implies that ipTM is at best a coarse proxy for specificity-relevant interface recovery across different models rather than a reliable discriminator on its own.

To assess the impact of data leakage, we examined epitope recall distributions for real complexes split into train-leakage and true test sets (Figure 4B). Across models, train-leakage complexes showed slightly higher median epitope recall than true test complexes, consistent with partial memorization or bias toward known interfaces. However, the overall distributions remained broad, and substantial overlap persisted between train-leakage and true test predictions. Notably, shuffled complexes spanned similar recall ranges, indicating that leakage alone does not explain the limited discriminatory power of epitope recall. Thus, high epitope recall does not reliably indicate cognate binding even in the absence of explicit train/test overlap.

We next tested whether increasing stochastic sampling improves epitope discrimination by assessing epitope enrichment significance as a function of replicate count (Figure 4C). Enrichment significance was computed using a one-sided binomial test for each VHH-antigen pair. Let *N* denote the number of stochastic structure predictions and *K* the number of predictions containing at least one epitope contact. Under a null model in which contacts occur uniformly across the antigen surface with probability *p_epitope_* (estimated from the fraction of solvent-accessible antigen residues (rSASA) belonging to the epitope), the probability of observing *K* or more epitope hits is given by the binomial survival function *P*(X ≥ K | X ∼ Binomial(N,p_epitope_)). Note that this is a deliberately coarse null. In practice, VHH–antigen contacts are spatially heterogeneous due to local geometry and physicochemical ‘hotspots’ (e.g., concavities, charge patches, domain accessibility), so a uniform-over-SASA model can miscalibrate absolute enrichment p-values. We therefore interpret these enrichment statistics primarily as an operational, antigen-normalized measure of epitope-seeking behavior, rather than a physically faithful generative null. Because the same null is applied to real and shuffled pairs for a given antigen, this limitation is less likely to explain the similarity of the saturation curves between real and shuffled cohorts, but it does limit interpretation of FDR as a calibrated significance level.

Resulting p-values were corrected across all VHH-antigen pairs using the Benjamini-Hochberg procedure, and pairs with FDR < 0.05 were considered significantly enriched. To assess the effect of ensemble size, we recomputed binomial p-values for hypothetical replicate counts *m* by rescaling the observed hit fraction (K/N) and repeating the same multiple-testing correction. This analysis also treats stochastic replicates as independent Bernoulli trials with a common hit probability. Diffusion replicates may exhibit non-trivial correlation because they share the same conditioning signal (sequence/MSA/embeddings) and differ only by sampling noise, reducing the effective sample size relative to N. As a result, p-values (and the extrapolation to larger m) may be optimistic, and therefore emphasize relative, within-model comparisons and the qualitative saturation behavior rather than interpreting FDR thresholds as strictly calibrated. For all three models, the fraction of VHH-antigen pairs declared significantly enriched increased rapidly with the number of stochastic predictions and was saturated by approximately 10-15 replicates. Importantly, real and shuffled complexes exhibited highly similar saturation behavior, regardless of whether the shuffled pairs involved train leakage of the VHH, the antigen, both, or neither. This indicates that increasing ensemble size primarily reinforces existing contact preferences rather than sharpening discrimination between cognate and non-cognate interactions. AF3 and Boltz-2 retained a modest residual separation between real and shuffled complexes across replicate counts, with real pairs showing slightly higher significant-enrichment rates than any shuffled cohort. In contrast, Chai-1 showed near-complete convergence: shuffled complexes rapidly achieved significance frequencies comparable to (or exceeding) those of real complexes, consistent with an epitope-seeking prior under this enrichment definition that limits the interpretability of enrichment-based metrics for this model. Notably, the leakage-stratified shuffled cohorts largely overlapped within each model, further indicating that these effects are not primarily driven by explicit train/test leakage but instead reflect intrinsic model priors that are amplified by ensembling.

Epitope recall of each model was then correlated against the ipTM values of the predicted, real, complexes. Figure S2B compares each model’s ipTM (diagonal) to system-level epitope-recall (diagonal). AF3 and Boltz-2 show weak association (pearson r≈0.18 and 0.08; spearman ρ≈0.15 and 0.14 respectively), whereas Chai-1 shows a moderate correlation (r≈0.46; ρ≈0.48). However, ipTM is designed as a global complex-confidence measure rather than an epitope-localization score, so high ipTM can occur even when the interface is confidently placed on the wrong antigen region. Moreover, ipTM is range-restricted for many systems (notably Boltz-2), and epitope-recall is bounded with a substantial mass near zero, making linear trendlines and correlation coefficients descriptive rather than calibrated. Additionally, since multiple pairs can share the same antigen (and/or related antigens), observations are not strictly independent and correlation magnitudes should be interpreted qualitatively.

Finally, we examined the relationship between structural consistency across predictions and epitope recovery within each model (Figure 4D). Structural consistency was quantified as the mean pairwise RMSD across n=50 stochastic replicates for each complex. Across all models and data splits, epitope recall was negatively correlated with replicate RMSD, indicating that complexes predicted with consistent orientations tend to recover a larger fraction of epitope residues. However, this relationship holds for both train-leakage and true test complexes, and does not differentiate real from shuffled pairings. pearson correlation coefficients ranged from r=-0.22 to -0.30 for AF3, r=-0.14 to -0.27 for Boltz-2, and r=-0.39 to -0.59 for Chai-1, demonstrating that while structural consistency improves epitope recovery, it does not guarantee biological correctness. It is worth noting that RMSD distributions for 50 replicas of real and shuffled complexes show slightly higher values for the shuffled complexes than for the real complexes (Figure 4E), suggesting a potential metric for distinguishing cognate and non-cognate interfaces.

Together, these analyses show that epitope recovery and enrichment metrics are driven primarily by generic contact biases and structural consistency rather than by interaction specificity. Increasing stochastic sampling improves apparent epitope enrichment but does so similarly for real and shuffled complexes, limiting its utility as a discriminative signal. These findings reinforce the conclusion that current structure prediction tools preferentially identify plausible antigen contact regions, but struggle to distinguish true cognate paratope-epitope interactions from non-cognate alternatives.

### Computational cost and sampling efficiency differ substantially across structure prediction tools

Using ML structure prediction tools for high-throughput antibody/nanobody screening requires understanding not only prediction accuracy but also computational cost. With GPU resources unevenly distributed across institutions ^47^ and growing concern over the energy footprint of large-scale computation ^48,49^, identifying cost-effective sampling strategies has practical importance. Saturation sampling and seed selection can both improve prediction quality ^15,26,31,50^, yet their relative contributions and associated costs remain uncharacterized. We therefore evaluated each system (n=106, 91 unique PDBs; no deposition-year bias observed; Supplementary Figure 6A) across five random independent seeds with increasing numbers of diffusion samples (N = 1, 10, 25, 50, and 100, around 100,000 additional structures in total), monitoring GPU energy consumption throughout (Figure 5). For this analysis, we additionally included Boltz-1, the predecessor to Boltz-2, to check the consistency of saturation effects across model versions. Because each saturation level uses an independent seed, this design enables analysis of combined seed and saturation effects; extracting only the first sample from each seed isolates seed-dependent variation.

**Figure 5.**
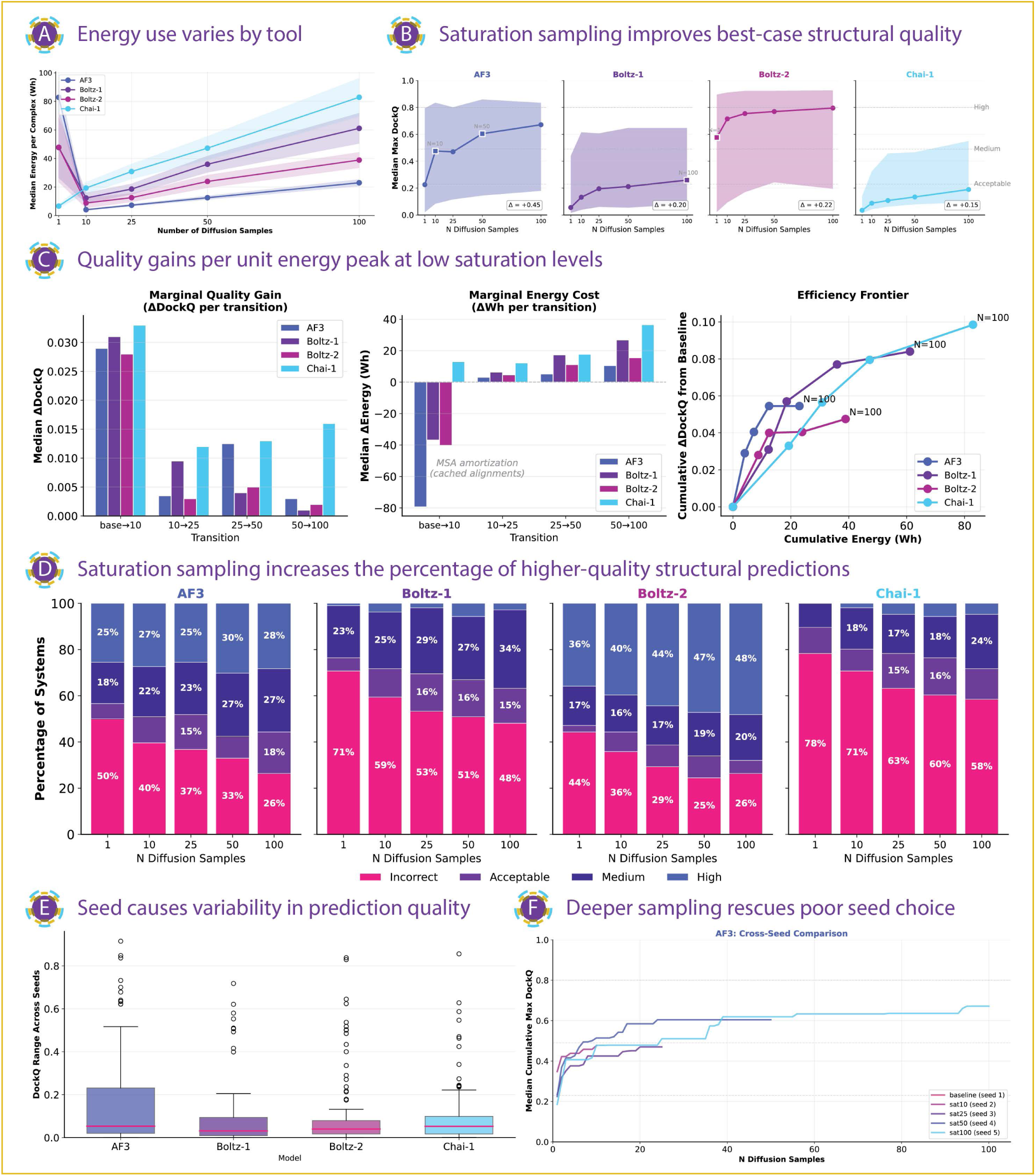
Computational cost, sampling efficiency, and seed-dependent optimization in nanobody-antigen structure prediction. A. *Energy consumption differs substantially across tools and sampling depth.* Median energy usage per predicted complex is shown as a function of diffusion sampling depth (N=1, 10, 25, 50, 100), with each saturation level corresponding to an independent random seed. Shaded regions indicate interquartile ranges across 106 systems. At the highest saturation level (N=100), median energy consumption spans from 23.0 Wh for AF3 to 82.9 Wh for Chai-1, reflecting distinct baseline and scaling behaviors across architectures. B. *Saturation sampling improves best-case structural quality with model-dependent magnitude.* Median maximum DockQ is shown as a function of sampling depth, with shaded regions indicating interquartile ranges across systems. Because each saturation level uses an independent seed, curves reflect combined effects of seed selection and increased sampling. Total improvements from baseline to N=100 (ΔDockQ) are indicated for each model, with AF3 showing the largest gain. Horizontal dashed lines mark DockQ quality thresholds separating incorrect (<0.23), acceptable (0.23-0.49), medium (0.49-0.80), and high (≥0.80) quality regimes. C. *Quality gains per unit energy peak at low saturation levels.* Bar plots show the median gain in maximum DockQ between consecutive saturation levels across systems. The largest improvements occur between baseline and low saturation, with progressively smaller gains at higher sampling depths. Each transition reflects both a change in seed and an increase in the number of samples. Line plot shows that efficiency frontiers relate cumulative DockQ improvement relative to baseline (seed 1, N=1) to cumulative energy expenditure. Each point corresponds to a saturation level and its associated seed. Steeper initial slopes indicate high computational efficiency at low sampling depths, followed by pronounced flattening at higher energy costs, consistent with diminishing returns beyond moderate saturation. D. *Saturation sampling reduces incorrect predictions but plateaus at higher depth.* Stacked bar plots show the distribution of DockQ quality categories across saturation levels, with each level corresponding to an independent seed and its associated sampling depth. The fraction of incorrect predictions (DockQ<0.23) decreases substantially at low saturation but shows limited further reduction at higher sampling depths. E. *Seed selection introduces substantial variability in prediction quality.* Boxplots show the range of DockQ values obtained from five independent seeds using only the first sample (sample₀), isolating seed effects from saturation. For each system, the DockQ range (maximum minus minimum across seeds) highlights that a subset of predictions exhibits large seed-dependent variation without additional computational cost. F. *Different seeds explore distinct solution landscapes.* Cross-seed saturation trajectories for AF3 show median cumulative maximum DockQ across systems as a function of sample number within each seed’s saturation run. Seeds that achieve higher quality early maintain their advantage throughout sampling, whereas poor initial seeds plateau at lower quality ceilings, indicating that seed selection and saturation sampling act as orthogonal optimization mechanisms. Corresponding analyses for other models are shown in Supplementary Figure 6.

To characterize how computational cost scales with sampling depth, we computed the median energy consumption across all systems at each saturation level (N = 1, 10, 25, 50, and 100). Energy consumption varied markedly between tools, revealing distinct cost profiles. At the highest saturation level (N=100 samples), median energy usage per system ranged from 23.0 Wh for AF3 to 82.9 Wh for Chai-1, corresponding to a 3.6-fold difference (Figure 5A, Supplementary Figure 5A-C). Boltz-1 and Boltz-2 exhibited intermediate energy-usage profiles, with substantially lower cost than Chai-1 at high saturation but higher baseline costs than AF3. When aggregated across all 106 real systems and saturation levels, total energy consumption ranged from 13.6 kWh for AF3 to 22.7 kWh for Chai-1.

These differences reflect different architectural choices. AF3 incurred a high baseline energy cost for MSA computation (82.4 Wh per system, computed once regardless of sampling depth), but showed relatively low marginal cost for additional diffusion samples. In contrast, Chai-1 exhibited a low baseline cost (6.6 Wh) but a steep increase in energy consumption with increasing saturation, consistent with its embedding-based workflow that omits MSA computation (Methods). Boltz-based models showed intermediate behavior. Across all tools, both energy consumption and time scaled approximately linearly with system length, with pearson correlation coefficients ranging from r=0.60 for AF3 to r=0.98 for Boltz-1 (Supplementary Figure 5D,E).

Increasing saturation sampling improved prediction quality for all models with diminishing returns, as measured by DockQ (Figure 5B). We quantified this by identifying, for each system, the maximum DockQ achieved at each saturation level, then computing the median across systems. Median maximum DockQ increased from baseline (N=1) to N=100 by Δ=+0.43 for AF3 (0.24 to 0.68), Δ=+0.23 for Boltz-2 (0.57 to 0.80), Δ=+0.20 for Boltz-1 (0.06 to 0.26), and Δ=+0.15 for Chai-1 (0.04 to 0.19). While all models benefited from increased sampling, the magnitude of improvement differed substantially, with AF3 showing the largest gains.

Marginal quality gain was computed as the per-system difference in maximum DockQ between consecutive saturation levels, then summarized as the median across systems. Across models, the largest marginal improvements occurred at low saturation levels. The transition from baseline to N=10 samples captured the majority of achievable quality improvement, while additional sampling beyond N=25 yielded progressively smaller gains (Figure 5C). This pattern is reflected in marginal ΔDockQ per sampling transition, which peaks at the base to 10 transitions across all tools and declines sharply thereafter. Consistent with this pattern, the proportion of incorrect predictions (DockQ<0.23) decreased sharply from baseline to N=10 but showed more modest (∼5-15%) reductions at higher saturation levels (N=25-100, Figure 5D).

Analyzing cumulative DockQ improvement as a function of cumulative energy expenditure revealed a consistent “efficiency frontier” across tools (Figure 5C). All models exhibited steep initial slopes, indicating high quality gains per unit energy at low sampling depths, followed by pronounced flattening as energy expenditure increased. Despite differing absolute costs, the efficiency frontiers converge across tools, indicating that early sampling dominates in terms of quality gains regardless of architecture. These trends indicate that N∼10-25 samples capture most of the attainable quality improvement at a fraction of the computational cost required for N=100, regardless of model architecture.

Beyond sampling depth, seed choice had a substantial independent effect on prediction outcomes (Figure 5E). To isolate seed effects from saturation, we extracted only the first sample (sample₀) from each seed and computed the per-system DockQ range (maximum minus minimum across seeds). The median range was 0.04-0.05, though extreme cases spanned nearly the full quality spectrum (e.g. system_70_70_8EN2: DockQ 0.01-0.93 across five different AF3 seeds). Across models, approximately 10-15% of systems were strongly seed-sensitive and benefited disproportionately from deeper saturation sampling (Supplementary Figure 6D, Supplementary Figure 7A).

Cross-seed trajectory analysis further showed that deeper sampling can partially rescue poor initial seed choices. For each seed, we tracked cumulative maximum DockQ (the best quality achieved up to each sample number) and computed the median across systems (Figure 5F). While seeds producing high-quality structures early tended to maintain their advantage throughout saturation, seeds that began in low-quality regions could still achieve substantial improvements with deeper sampling, though typically plateauing at lower absolute quality than favorable seeds (Figure 5F, Supplementary Figure 7B). Together, these results indicate that seed selection and saturation sampling act as orthogonal optimization mechanisms: seed choice determines which region of the solution landscape is explored, while saturation sampling refines predictions within that region and can partially mitigate but rarely fully overcome suboptimal initial trajectories (∼85-90% of systems remained below their best-seed ceiling).

Finally, we assessed whether confidence metrics reflect structural improvements achieved through saturation sampling. Across all models, changes in ipTM from baseline to N = 100 showed near-zero correlation with corresponding changes in DockQ (pearson r=0.00-0.16; Supplementary Figure 6B,C). This indicates that confidence scores are largely determined by properties of the initial prediction and remain insensitive to substantial gains in quality from additional sampling. Reliable confidence tracking would enable early stopping without ground-truth structures, but the observed ipTM-DockQ decoupling precludes such use. This behavior mirrors the DockQ-ipTM misalignment observed above, reinforcing that current confidence metrics do not reflect convergence toward higher-quality docking solutions.

Together, these results demonstrate that moderate sampling (N∼10-25) captures most achievable quality gains at a fraction of full saturation cost, seed selection and saturation act as orthogonal optimization mechanisms, and current model confidence metrics fail to track these improvements for VHH-antigen predictions.

## Discussion

This work benchmarks whether modern AI complex-prediction tools can discriminate cognate nanobody-antigen binding and recover correct paratope-epitope interfaces. Our central findings are: across tools, high-level confidence scores frequently fail to separate real from mismatched (“shuffled”) complexes, and sampling can improve best-case geometry without resolving the upstream problem of selecting the correct binding mode. Importantly, even when structural quality improves substantially under aggressive sampling (as, for example, measured by DockQ for real complexes) model confidence scores often remain unchanged, revealing a disconnect between structural refinement and confidence calibration. Below, we place these results in the broader “AI biologics” landscape and outline practical implications for drug discovery and model development.

### Where the field stands: disconnect between de novo generation and de novo prediction

The AI biologics ecosystem is now shaped by strong competition among major players (e.g., Latent, Chai, Nabla, Isomorphic, Boltz), with rapid iteration cycles. Although de novo generation success rates are increasing into the double digit realm (∼10-15% on hard interface tasks ^30,51^), our results indicate that there is a disconnect between de novo design (task: “generate an antibody for a given target without exploiting other information”) and de novo prediction (task: “indicate all the antigens that can be bound by a given antibody, and vice-versa”). Our “real vs shuffled” discrimination benchmark shows that models can generate interfaces that appear plausible, albeit incorrect, across many pairings, resulting in an abundance of false positives that are challenging to triage.

An emerging industry screening workflow is: generate thousands to millions of candidates, then filter using internal confidence scores (e.g., pLDDT, ipTM, PAE). Our results suggest that this strategy is fragile for antibody-antigen binding, because confidence does not necessarily equate to accurate biology. In our many-VHH-versus-many-antigen screening setting, shuffled complexes often reach confidence levels comparable to cognate pairs, while genuinely high-quality structures can remain underconfident depending on the prediction tool. This breaks the assumption that “high-confidence docking implies correct binding” and it helps explain why confidence-guided selection can produce many experimental failures even when predicted structures look polished. Consistent with recent proposals for alternative evaluation metrics ^52,53^ our results underscore the need for “better scores” that more directly reflect biological meaning rather than internal structural self-consistency.

Crucially, drug viability depends on properties that these scores do not directly encode: induced fit compatibility, entropic penalties, off-target propensity, developability constraints, and the ability to tolerate antigen dynamics or conformational selection ^54,55^. Confidence metrics were not designed as surrogates for molecular functionality, and our data reinforce (with regard to off-target binding potential, for example) that treating them as such inflates false positives, especially in interface-dependent problems.

### Why evolutionary signals and sampling help less than hoped for antibodies

A common intuition is that better MSAs and more evolutionary information should improve AI docking and design. However, both antibody and nanobody paratopes (especially CDR loops) represent the most variable regions in biology, and much of the binding specificity arises from flexibility, conformational diversity, and context-dependent loop rearrangements rather than sequence conservation ^56–58^. This is consistent with the idea that the utility of MSAs in antibody-focused tasks is more nuanced than in general protein structure prediction. While MSAs are clearly critical for accurate monomer modeling (removing them in AF3 leads to large degradations in structural accuracy ^59^), their contribution to antibody-antigen docking is constrained by the high sequence and conformational diversity of CDRs, which limits the availability of informative evolutionary signals. This suggests that MSA-derived confidence estimates may be inherently less reliable for antibody-antigen interfaces than for other protein complexes ^60^. As a result, alternative MSA-strategies could provide better guidance; especially in cases of evolutionary complex/highly mutating proteins such as antibodies.

On the other side, sampling helps, but exposes a deeper bottleneck: path selection vs refinement. Saturation/diffusion sampling improved best-case DockQ in many systems, confirming that sampling is valuable as a refinement mechanism. However, we observe essentially zero correlation between changes in DockQ (ΔDockQ) and changes in ipTM (ΔipTM) across all three models (pearson’s r=-0.03, -0.04, and -0.02 for AF3, Boltz-2 and Chai-1), indicating that confidence scores do not track actual structural improvement. The persistence of “non-improver” systems supports a two-stage view: i) path/seed selection determines the qualitative docking mode (often wrong) and ii) saturation refines within that chosen mode.

In this sense, current models appear to have a “fixed mindset”: once a confidence level is assigned to a given seed or trajectory, it remains largely invariant, even when the resulting structure improves substantially. This likely reflects an architectural constraint: confidence scores (like ipTM) are derived from MSA-based pairwise representations computed in the trunk network, which remains fixed during diffusion sampling ^61^. Indeed, MSA representations can be directly optimized via gradient descent to manipulate confidence outputs ^61,62^, confirming this upstream dependency. If the model commits early to an incorrect binding mode, additional diffusion samples often cannot rescue it. This has immediate practical implications: “just sample more” may be an expensive way to get diminishing returns, and large seed sweeps, while sometimes recommended, can be prohibitive at screening scale-without guaranteeing specificity or providing confidence-based validation that sampling actually helped. Development of confidence measures sensitive to binding mode quality-or orthogonal strategies such as consensus across independent seeds-remains an important challenge for the field.

### Structural biology perspective: flexibility is not optional for paratope-epitope recovery

Antibody recognition is not merely geometric matching; it is often a dynamic process where CDR loops adapt to the antigen surface and exploit transient conformations ^63^. This is particularly relevant for targets with disordered regions or multiple accessible states. Therefore, a major unresolved challenge is distinguishing hallucinated (geometrically plausible but physically or functionally implausible) complexes from viable ones that can maintain binding under realistic dynamics.

This points to a missing ingredient in many AI pipelines: a standardized representation of flexibility and experimentally grounded dynamic behavior. We argue that molecular dynamics (MD)-informed filtering is a promising bridge between AI-generated hypotheses and functional plausibility ^4,64,65^. However, today’s AI-generated “MD-like” outputs are often not physically faithful enough to serve as ground truth. These insights suggest that future antibody-antigen discovery pipelines should move beyond static confidence metrics toward standardized, dynamics-aware frameworks, such as MD-informed benchmarks and datasets like DINO ^65^, that prioritize biophysical plausibility, interface stability, and functional robustness to enable efficient selection of truly viable therapeutic candidates.

### Limitations and outlook

Our study focuses on VHH-antigen complexes under a controlled real-versus-shuffled pairing design, which is intentionally challenging: it tests specificity (off-diagonal), not just docking plausibility (on-diagonal). In other words, our benchmark quantifies the extent to which generalized (or universal) binding prediction is feasible. That said, our study may overestimate performance limitations in narrower settings where the antigen epitope is known, constraints are available, or the target is rigid and structured. Furthermore, some of the assumed non-binding shuffled complexes may indeed be binding (off-target effect is typically less than 1% ^66^). Additionally, similar benchmarking should be extended to more complex antibody formats to assess whether these findings generalize beyond VHHs.

Recent large-scale resources such as AbSet ^53^ (a dataset of over 800,000 antibody structures including both experimental and computational models) highlight an additional challenge: the sheer volume of predicted antibody structures does not imply reliability. Systematic assessment of how many computational antibody structures in such datasets are structurally sound, interface-correct, or functionally meaningful remains limited. Our results are consistent with the view that antibody structure prediction, particularly in complex with antigens, is not yet a solved problem. Quantifying the extent to which prediction performance is a function of structural data quality in train and test sets remains to be investigated.

Another important limitation concerns the structural evaluation metrics employed in this study. While DockQ provides an established composite measure of interface quality, it remains a geometry-centric metric and does not directly assess specificity, energetic plausibility, or functional robustness. Future work should therefore systematically investigate alternative and complementary interface-focused measures ^36^, such as ipSAE ^61^, FNAT (fraction of native contacts) ^67^, CAPRI-style classifications ^68^, interface RMSD variants ^69^, and other contact-overlap ^56^ or energy-informed metrics ^70–74^, to determine whether they better correlate with cognate discrimination and biological plausibility than trunk-derived confidence scores (e.g., ipTM). A broader metric landscape may reveal evaluation signals that are more sensitive to binding mode correctness or to subtle interface rearrangements that are not captured by current confidence outputs.

Looking forward, progress will likely require: (1) interface-specific training objectives that incorporate hard mutation-diverse negatives (shuffled non-cognate pairings) across several affinity ranges ^75–79^, (2) confidence estimates calibrated to specificity and functional plausibility rather than structural self-consistency, and (3) standardized dynamic datasets ^65^ enabling models and filters to account for flexibility and induced fit. The most impactful near-term improvement may not be a new confidence score, but a robust, scalable post-prediction filtering layer grounded in biophysics and dynamics.

### Guidance on sampling, compute-accuracy tradeoffs, and confidence calibration

Our results have immediate consequences for how AI-based structure prediction is operationalized in antibody and antibody discovery pipelines. In current industry practice, large libraries of candidates are frequently filtered using internal confidence scores (e.g., ipTM, PAE, or pLDDT), often combined with extensive stochastic sampling to improve predicted geometry. However, the analyses presented here indicate that such workflows risk conflating structural plausibility with binding specificity. Because shuffled non-cognate complexes frequently achieve confidence scores comparable to real complexes, and because confidence remains largely insensitive to structural improvements achieved through sampling, we argue that future screening strategies must explicitly decouple geometric refinement from interaction prioritization.

A key practical implication concerns the role of stochastic sampling. Across all models, moderate sampling improved best-case structural quality, confirming that diffusion sampling is valuable as a refinement mechanism. Yet additional samples primarily refine an already selected structural hypothesis rather than enabling discrimination between cognate and non-cognate partners. Note that the reported gains reflect best-of-ensemble outcomes (i.e., selecting the highest-quality structure among samples using ground-truth evaluation), which is not directly available in prospective pipelines. Deeper sampling only improves decision-making to the extent that downstream filters can reliably identify the better mode among the sampled structures; otherwise, additional samples mainly increase the number of plausible-looking false positives. In practice, this suggests that sampling should be used as a mode-exploration step rather than as a simple confidence amplifier. For high-throughput workflows, shallow ensembles (≈10-25 stochastic predictions) capture most achievable DockQ improvement at a fraction of the computational cost of deep saturation runs, while larger ensembles mainly yield diminishing returns. Importantly, seed choice and sampling depth act as orthogonal optimization mechanisms: independent seeds explore different regions of the solution landscape, whereas deeper sampling refines within those regions. Consequently, distributing computational budget across multiple seeds is often more informative than increasing sampling depth within a single trajectory. Treating sampling as a tool to detect structural convergence rather than as a guarantee of specificity may reduce unnecessary GPU expenditure without compromising structural insight.

These observations also inform compute-accuracy tradeoffs in large-scale biologics programs. The efficiency frontiers observed across tools show steep early gains in quality per unit energy, followed by rapid flattening at higher sampling depths. From a discovery perspective, this implies that the most cost-effective strategy is a staged workflow: early, low-cost ensembles to identify stable interface modes, followed by selective deeper sampling only for candidates that satisfy additional biological or experimental constraints. Because confidence scores do not track structural refinement, escalating compute solely to improve ipTM values is unlikely to yield more reliable candidate prioritization. Instead, compute allocation should be conditioned on whether additional sampling changes the structural hypothesis (for example, by resolving competing interface clusters or rescuing seed-sensitive systems) rather than on absolute confidence thresholds.

Perhaps the most consequential implication relates to confidence calibration. Our benchmark demonstrates that ipTM scores cannot be interpreted as probabilities of correct binding or as reliable indicators of specificity in realistic discovery settings. More precisely, our analyses show that these scores are not calibrated to specificity (cognate vs non-cognate binding) in this discovery-like setting. Converting ipTM (or any internal score) into a probability of correct binding would require target- and program-specific calibration data with explicit negatives and, ideally, affinity/function labels; such a mapping is not identified by structure-only benchmarking. Any learned calibration is also likely to drift across model releases and prompting/inference settings, motivating routine re-calibration within each program. Overconfident failures, underconfident successes, and weak cross-tool agreement all highlight that current confidence metrics capture internal structural consistency rather than biological correctness ^80^. For pharma-facing workflows, this necessitates a shift from absolute confidence thresholds toward relative or context-aware calibration. One practical approach is to compare each candidate against a panel of negative controls (such as shuffled pairings, homologous off-targets, or unrelated antigens) and to evaluate whether predicted interfaces are exceptional relative to these decoys. Under such a framework, confidence becomes a comparative signal embedded within a program-specific landscape rather than a universal scalar metric. This paradigm aligns with our real-versus-shuffled benchmarking strategy and reflects the reality that specificity can only be assessed in the presence of realistic alternatives.

Beyond score calibration, ensemble-derived signals may provide a more robust basis for decision-making. Structural consistency across stochastic predictions, dominance of a single interface cluster, and reproducibility of epitope contacts emerge as practical indicators of hypothesis stability, even though they do not by themselves guarantee biological correctness. Incorporating these ensemble-level features into downstream filtering layers may help distinguish fragile, hallucinated complexes from hypotheses worthy of experimental follow-up. Notably, cross-tool agreement on interface mode rather than agreement on raw confidence magnitude may represent a stronger indicator of robustness, given the low correlation in ipTM landscapes observed here.

Taken together, these considerations suggest a reframing of AI-assisted antibody discovery workflows. Rather than relying on confidence scores as standalone decision metrics, future pipelines may benefit from explicitly separating three conceptual layers: (i) geometry confidence, describing structural self-consistency, (ii) mode confidence, reflecting ensemble convergence toward a stable interface hypothesis, and (iii) specificity confidence, derived from comparative evaluation against realistic decoys or alternative partners. Such a multi-layered framework acknowledges the limitations identified in this study while preserving the strengths of modern structure prediction tools as generators of plausible structural hypotheses. In the near term, the most impactful improvements may therefore arise not from deeper sampling or higher raw confidence scores, but from better-calibrated filtering strategies that integrate ensemble behavior, negative controls, and biological context into a unified decision process for AI-guided biologics development.

## Methods

### VHH-antigen dataset curation

For benchmarking nanobody-antigen complex prediction, VHH-antigen structures were curated from two complementary sources: SAbDab-nano ^81^, downloaded March 2025, 90% sequence redundancy cutoff, bound complexes only) and the Antigen-nanobody Complex Database (AACDB) ^82^. To minimize overlap with the training data of AlphaFold 3 (AF3), Boltz-2, and Chai-1, only post-October 2021depositions were retained from SAbDab-nano, corresponding to the earliest training cutoff among the evaluated tools (Boltz-2). To explicitly probe potential memorization effects, pre-cutoff structures from AACDB were retained and later used to define train-leakage subsets.

All structures were filtered to include protein antigens only, with crystallographic resolution ≤ 3.0 Å, VHH length between 110-150 amino acids, and antigen length between 100-400 amino acids. The two datasets were merged and exact PDB-chain duplicates were removed. For epitope-paratope variation analysis, we defined a set of distinct VHHs binding the same antigen within a single PDB entry (Supplementary Table 1). This yielded 15 PDBs containing such replicates (30 systems).

To maintain computational feasibility, the total number of unique systems was capped below 125. After manual inspection, 17 systems were excluded due to structural artifacts, VHHs contacting multiple antigen chains, or ambiguous chain annotation. The final benchmark comprised 106 VHH-antigen systems from 91 unique PDB entries.

Antigen secondary structure was assigned using DSSP, amino-acid composition was computed directly from sequences, and PDB novelty was inferred from the first character of the PDB accession code.

### Real vs shuffled complex generation

To assess interaction specificity, we constructed a full combinatorial pairing matrix between the curated VHHs and antigens. For each VHH, the experimentally observed pairing with its cognate antigen was designated as the real complex, while all other VHH-antigen pairings were designated as shuffled complexes. This design preserves realistic molecular interfaces while systematically breaking biological specificity.

Unless otherwise stated, all analyses were performed on the complete 106 × 106 VHH-antigen matrix, enabling direct comparison between real and shuffled predictions under identical modeling conditions.

### Structure prediction

Structure predictions for the full combinatorial dataset (106 VHHs × 106 antigens, 50 samples per complex) were performed using three independent structure prediction frameworks: AlphaFold3 (AF3), Boltz-2, and Chai-1. Predictions were executed across multiple high-performance computing environments:

- AF3: Leonardo Booster partition (CINECA, Italy)
- Boltz-2: VEGA supercomputer (IZUM, Slovenia/University of Oslo)
- Chai-1: Immunohub cluster (University of Oslo)

Exact hardware specifications, runtime stacks, and software versions were logged at job start for reproducibility.

#### Chai-1

Chai-1 predictions were performed using version 0.6.1. All complexes were run on the University of Oslo Immunohub cluster using 1 GPU per job. Each system was predicted with 50 models, using the parameters: num_trunk_samples = 5, num_diffn_samples = 10, num_trunk_recycles = 3, num_diffn_timesteps = 200, seed = 42. No MSAs were computed; instead, ESM-based embeddings were used under Chai-1’s default inference mode

#### Boltz-2

Predictions were performed using the Boltz CLI v2.2.0. Boltz-1 and Boltz-2 were selected via --model boltz1 or --model boltz2, respectively. Unless otherwise noted, all runs used: --use_msa_server, recycling_steps = 3, sampling_steps = 200, diffusion_samples = 50, fixed random seed (42). Outputs were written in PDB or mmCIF format, with full per-residue and interaction-level scores exported using --write_full_pde. Boltz-2 jobs ran on the Immunohub and Boltz-1 on the VEGA.

#### AlphaFold3

AlphaFold 3 predictions were performed using the alphafold3-3.0.1 via Singularity container on the Leonardo Booster partition of CINECA. Each system was predicted with 50 diffusion samples using parameters: `num_diffusion_samples = 50`, `run_data_pipeline = true`, `run_inference = true`. Each job was run with a fixed random seed (`seed = 1`). MSAs were computed using the built-in data pipeline (`--run_data_pipeline=true`).

### Train, test, and mixed (“both”) labeling

Each VHH-antigen system was assigned to train, test, or both categories based on the training cutoffs of the corresponding model: i) AF3: ∼30 September 2021, ii) Chai-1: ∼12 January 2021, iii) Boltz-2: ∼1 June 2023. A system was labeled train if both the VHH and antigen were present in the model’s training set, test if neither was present, and both if one component was present while the other was not.

### Structural similarity between replicas

To quantify structural reproducibility, we computed geometric and contact-based similarity metrics across independently generated predictions. All analyses were performed on Cα atomic coordinates extracted from PDB files using custom Python scripts (NumPy ^83^, SciPy ^84^, pandas ^85,86^). For each complex, all pairwise comparisons among replicate structures were computed and aggregated to yield per-system metrics.

#### Root Mean Square Deviation (RMSD)

Global structural similarity was quantified using RMSD after superposition on the antigen via the Kabsch algorithm:

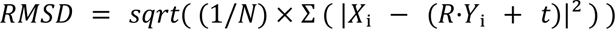

where *X* and *Y* are corresponding Cα coordinates, and *R, t* are the optimal rotation and translation. Lower RMSD values indicate higher structural consistency. For each system, RMSD values were averaged across all replicate pairs to obtain mean ± SD estimates.

#### Epitope enrichment

We computed epitope enrichment for each VHH-antigen pair using a one-tailed binomial test. For each complex (real or shuffled), a replicate prediction was scored as a hit if any contacted antigen residue overlapped the experimental epitope for that antigen. For each pair, we aggregated K (number of hit replicates) out of N (50 replicates). We assessed statistical enrichment using a one-sided binomial test under a null model in which contacts are uniformly distributed over the antigen surface. The null epitope-contact probability p_epitope_ was estimated per antigen structure as the fraction of antigen solvent-accessible surface area (SASA) attributable to epitope residues: p_epitope_ = SASA_epitope_/SASA_antigen_. Associated summary statistics including expected hits (N_pepitope_), enrichment lift ((K/N)/p_epitope_), and −log_10_(p). To control for multiple testing, p-values were corrected across all evaluated VHH-antigen pairs using the Benjamini-Hochberg procedure to obtain FDR-adjusted q-values.

To separate effects driven by cognate pairing versus dataset split leakage, each VHH-antigen pair was assigned a cohort label based on whether the pairing was real (cognate) or shuffled (non-cognate) and on the train/test split assignments of the individual binding partners.

Replicate-sensitivity (“significance vs. replicates”) analysis (Figure 4C) set out to evaluate how apparent enrichment depends on ensemble size; we performed a replicate-scaling analysis for hypothetical replicate counts *m*=1,…,N. For each pair, we approximated the number of hits at depth *m* by rescaling the observed hit fraction: Km=round((K/N) *m*). We recomputed one-sided binomial p-values using *m* and the same pepitope, applied Benjamini-Hochberg FDR correction across all pairs separately at each *m*, and recorded the fraction of pairs deemed significant (q<0.05) within each shuffled cohort. These cohort-stratified curves were used to summarize how quickly significance accumulates with increasing stochastic sampling for each model.

### Saturation analysis and computational cost measurement

To assess the relationship between computational cost and structural prediction quality, we performed saturation runs on VEGA (IZUM, Slovenia), with each job allocated a single NVIDIA A100 GPU (40 GB).

#### Sampling configurations

We tested diffusion sample counts of {n= 1 (baseline), 10, 25, 50, 100} for AlphaFold3 and Boltz-1/2. For Chai-1, we additionally varied trunk samples, testing configurations of {5×1, 5×2, 5×5, 5×10, 5×20} (trunk x diffusion samples), to match the structure prediction of the whole 106×106 matrix (as described above).

#### MSA strategy

To isolate inference costs from alignment overhead: i) AlphaFold3: MSAs were generated once per system during baseline run (diffusion_samples =1) and reused for subsequent runs via --run_data_pipeline=false, ii) Boltz-1/2: a shared MSA-cache (--cache) was populated during baseline runs and reused across all saturation runs, iii) Chai-1: ESM embeddings were used without MSA search (--use-ESM-embeddings)

#### Seed strategy

Each prediction job used independent random seed initialization to ensure independent sampling across runs. AlphaFold3 and Chai-1 used explicitly generated seeds at submission time; for Boltz-1/2 we used default random initialization. Within each run, multiple diffusion samples explore different conformations along independent stochastic trajectories.

#### Energy monitoring

GPU telemetry (power draw, utilization, memory usage) was sampled every 5 seconds using nvidia-smi and written to per-run CSV files. Wall-clock timing and system metrics were captured via /usr/bin/time -v. Total energy consumption (Wh) was calculated by trapezoidal integration of instantaneous power readings over the run duration:

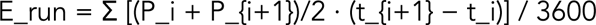

Runs were aggregated by (model × diffusion sample count) and reported as mean ± SD. Due to scheduling, initialization, and I/O overhead, reported values reflect relative efficiency under identical cluster conditions, not theoretical lower bounds.

### Data analysis and statistical aggregation

All analyses were implemented in Python using the NumPy, SciPy, matplotlib and pandas libraries. Pairwise structure comparisons were parallelized using the multiprocessing module. The workflow automatically parsed PDB files, performed optimal structural alignments, computed the defined similarity metrics, and aggregated results per system. For each complex system, all pairwise metric values were averaged to produce mean and standard deviation estimates.

#### Computational cost and sampling efficiency analysis

As mentioned above, energy consumption was measured using NVIDIA’s nvidia-smi tool, sampling GPU power draws at 5-second intervals throughout each prediction run.

Structural quality was assessed using DockQ v[2.1.3]^41^, which integrates interface RMSD, ligand RMSD, and fraction of native contacts into a single score ranging from 0 (incorrect) to 1 (perfect). Quality categories follow CAPRI conventions: incorrect (<0.23), acceptable (0.23-0.49), medium (0.49-0.80), and high (≥0.80).

For saturation analysis, each system was evaluated with five independent random seeds, each assigned to a single saturation level (N = 1, 10, 25, 50, and 100 diffusion samples for seeds 1-5, respectively). To isolate seed-dependent variation from saturation effects, we extracted only the first sample (sample_0_) from each saturation run and computed per-system DockQ and ipTM ranges (maximum minus minimum across five seeds). Chai-1 uses a fixed five-trunk architecture. Requiring a modified sampling scheme to achieve equivalent saturation levels. We varied the number of diffusion samples per trunk (1, 2, 5, 10 and 20 samples x 5 trunks = N of 5, 10, 25, 50, and 100 total samples). For seed-isolation analysis, we extracted sample_0_ from trunk_0_ as the baseline prediction for each seed.

Marginal quality gains were computed as the per-system difference in maximum DockQ between consecutive saturation levels, then summarized as the median across all systems. Efficiency frontiers were done by plotting cumulative median ΔDockQ (relative to baseline) against cumulative median energy expenditure at each saturation level. Cross-seed trajectory analysis tracked the cumulative maximum DockQ (i.e. the best DockQ achieved up to and including each sample number) within each seed’s saturation run, then computed the median across systems. Correlations between confidence score changes (ΔipTM) and quality changes (ΔDockQ) were assessed using pearson correlation coefficients.

## Data availability

Supplementary tables, experimental and computational data, along with related scripts and pipelines, can be found on Zenodo (10.5281/zenodo.18390239) and GitHub (https://github.com/csi-greifflab/ab_ag_champloo).

## Author contributions

E.S. and V.G. conceived the study. E.S. led the study, performed structure prediction using Chai-1 and Boltz-2, conducted data analysis, prepared figures, and wrote the manuscript. M.A. performed data analysis, prepared figures, and contributed to manuscript writing. K.K.B. performed structure prediction using AF3, conducted saturation and calibration studies, performed data analysis, prepared figures, and contributed to manuscript writing. L.S. and S.M. supported AF3 and Boltz-1 computations, including optimization and execution of prediction runs. A.K. and C.F. performed

sequence analyses and generated preliminary results. P.S. and A.d.M. supervised the study and provided scientific guidance. V.G. supervised the project, contributed to conceptualization, and participated in manuscript writing. All authors reviewed and approved the final version of the manuscript.

## Acknowledgements

We thank Prof. Charlotte Deane and Henriette Capel (OPIG, University of Oxford) for valuable discussions. We are grateful to Žiga Zebec (IZUM, Slovenia) for guidance on assessing VEGA and EuroHPC resources.

## Disclaimer

Co-funded by the European Union. Views and opinions expressed are however those of the author(s) only and do not necessarily reflect those of the European Union or European Research Executive Agency. Neither the European Union nor the granting authority can be held responsible for them.

## Supplementary material

**Supplementary Table 1.**
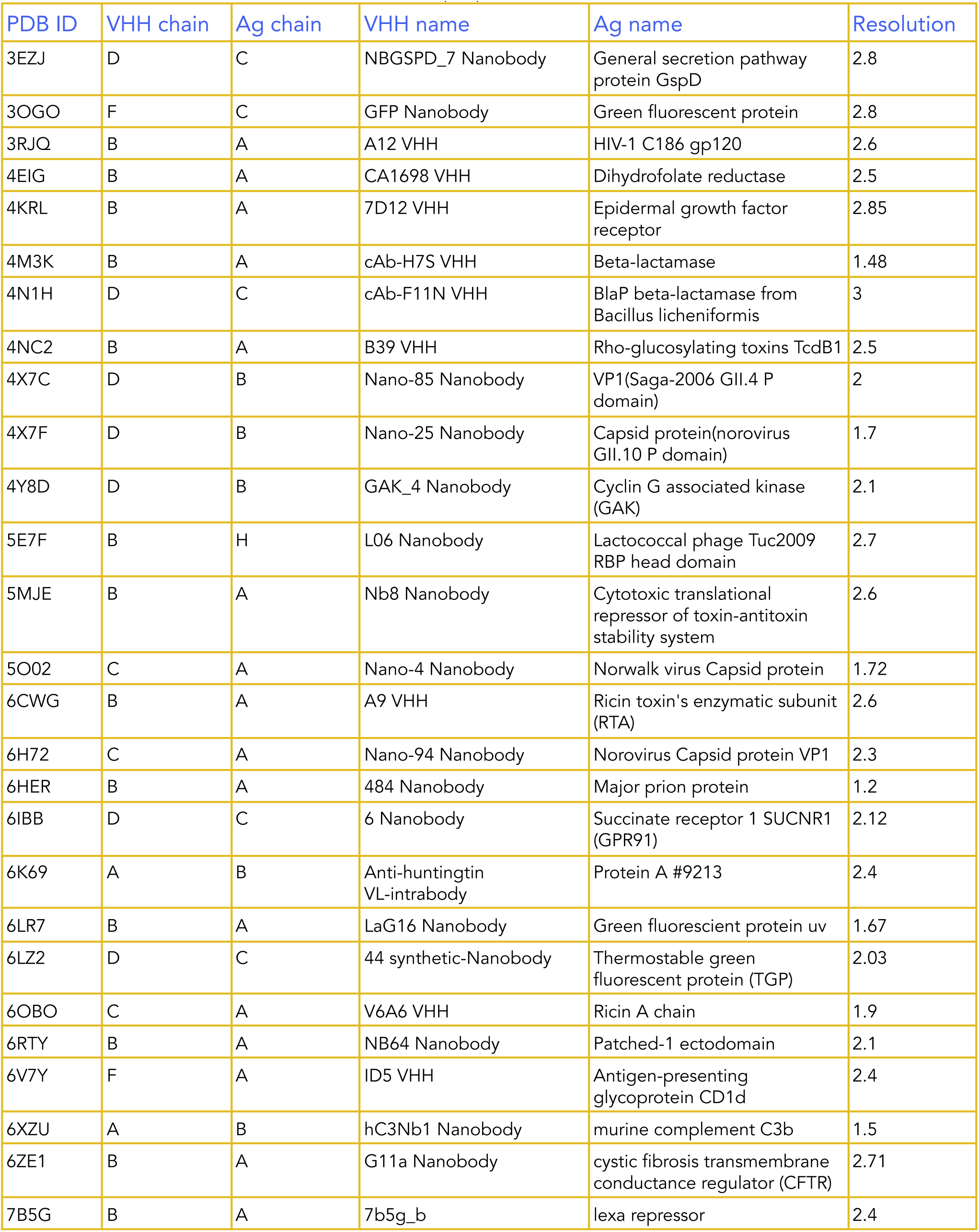
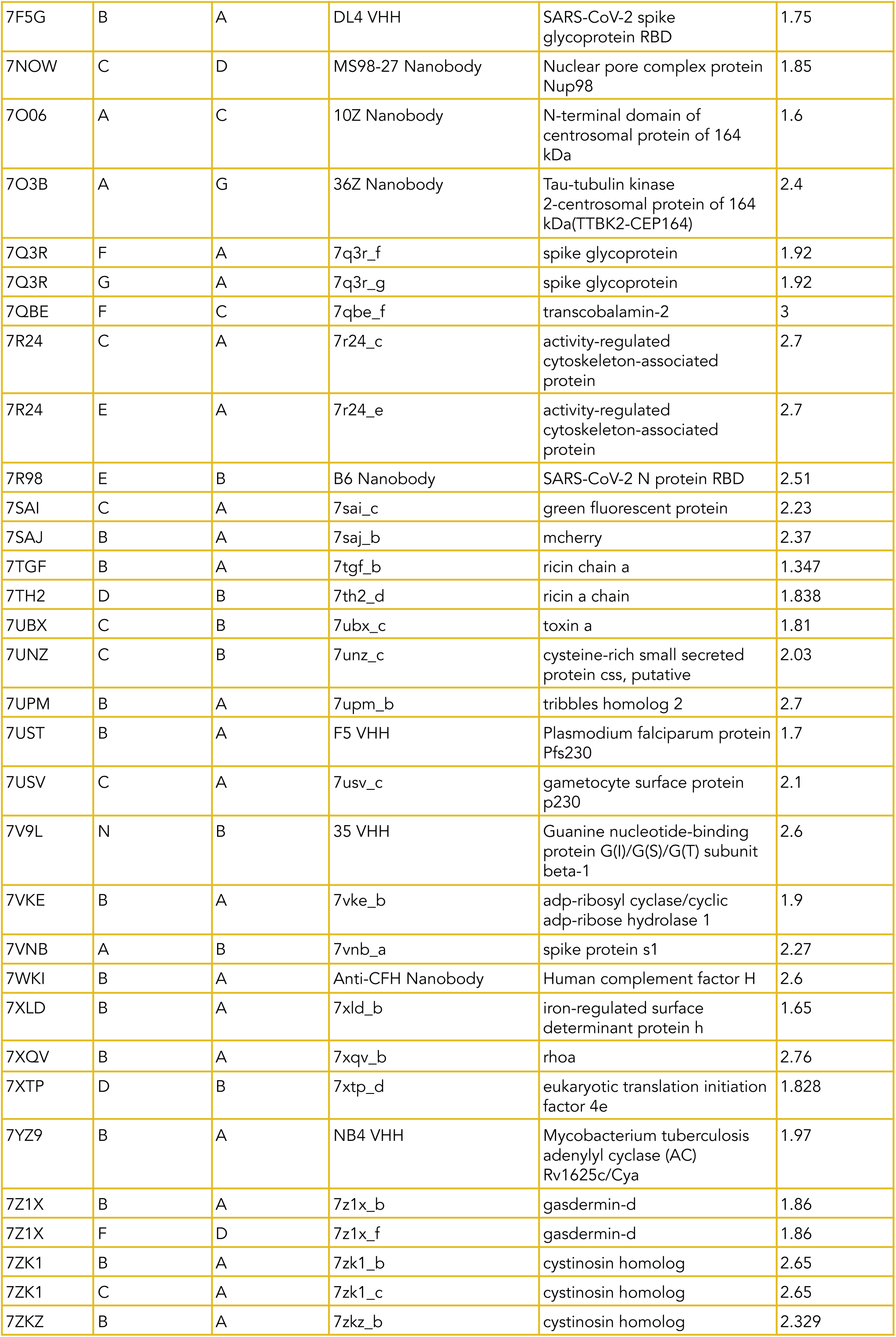
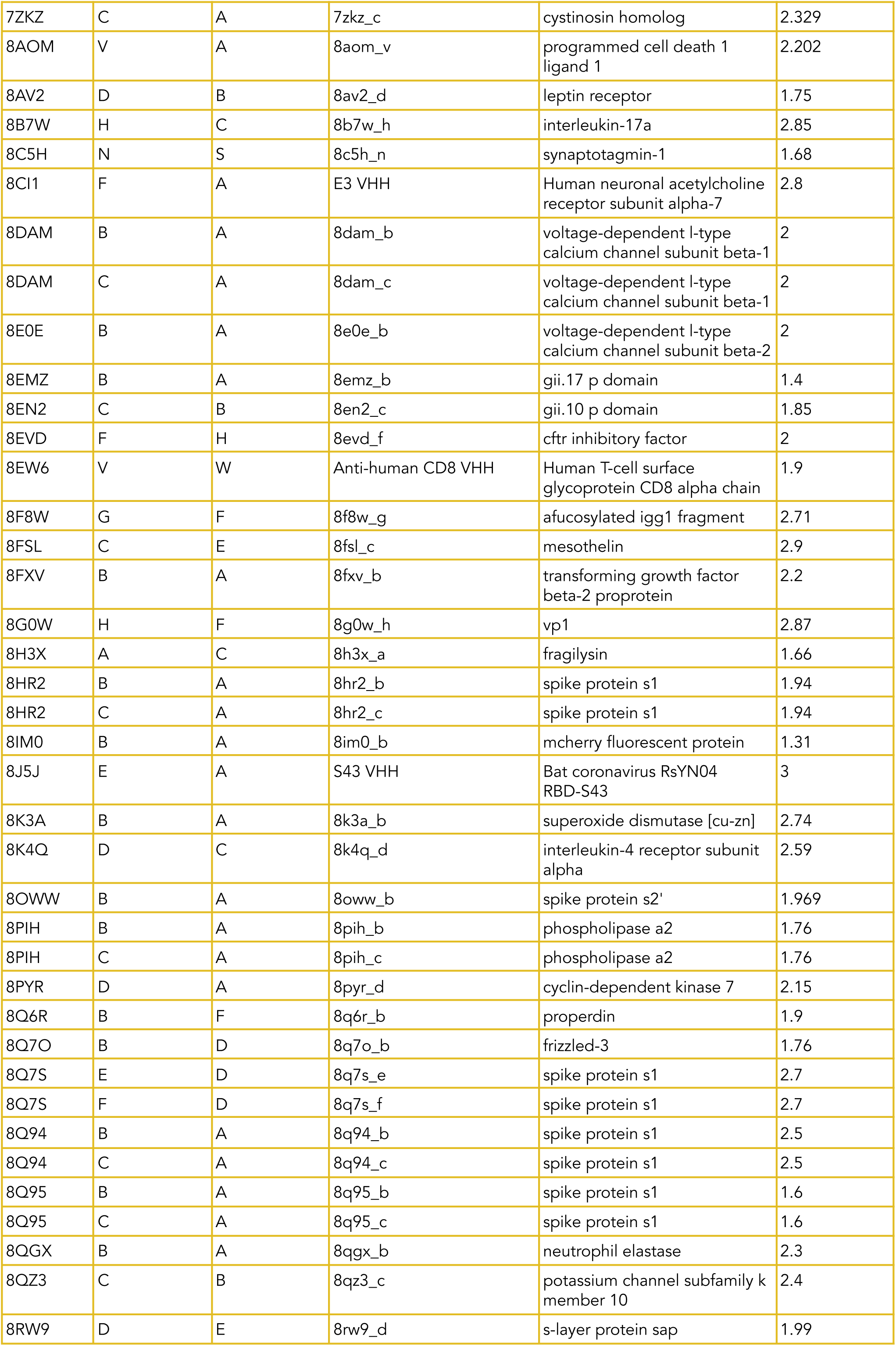
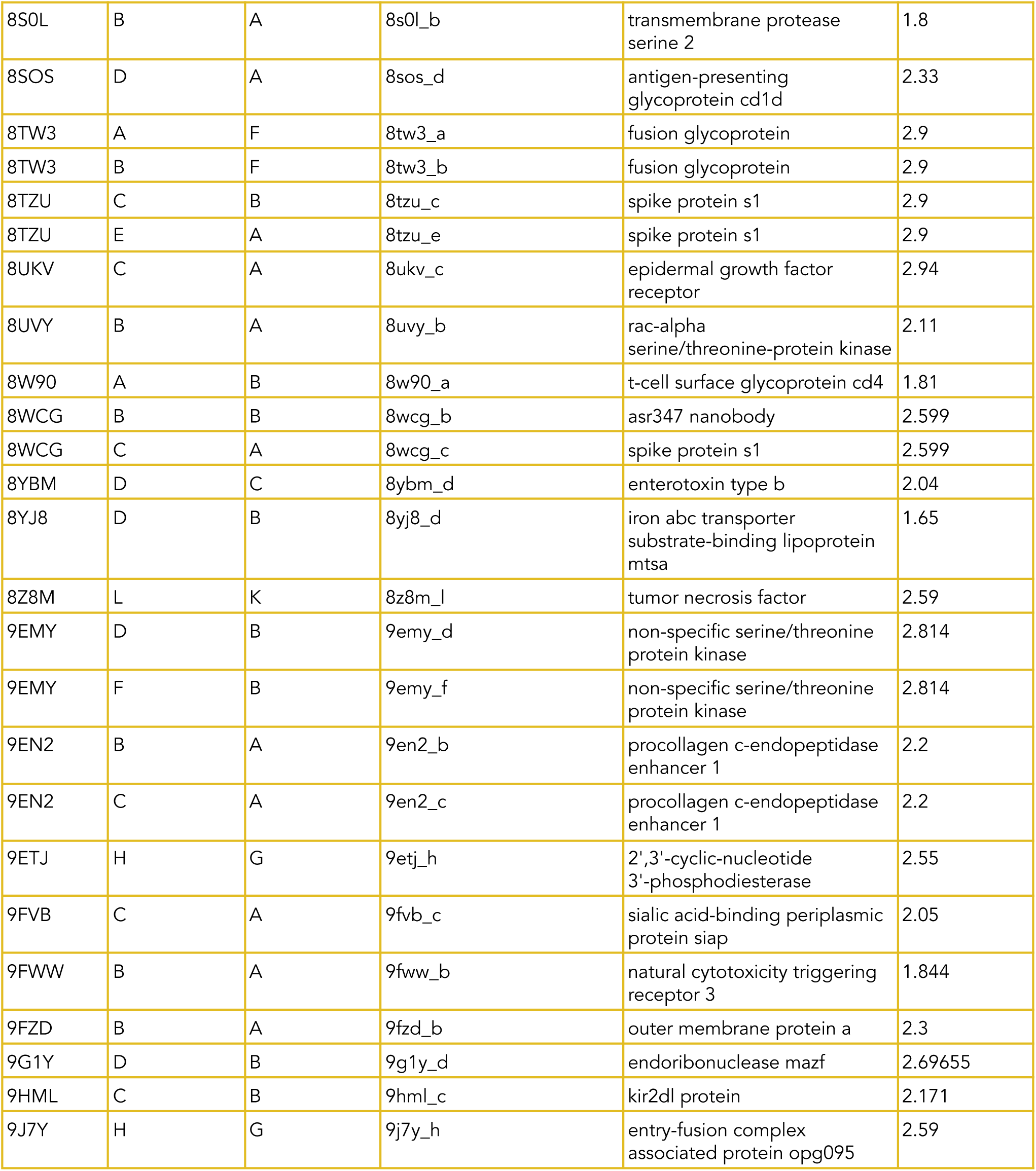
List of PDBs and the corresponding VHH and Ag chains from the crystal structures used in this benchmark. The table lists all Protein Data Bank (PDB) entries included in the benchmark, together with the specific VHH (nanobody) and antigen chain identifiers extracted from each crystal structure. For each complex, the corresponding nanobody name, antigen annotation, and experimental crystallographic resolution (Å) are provided. These structures form the ground-truth cognate complexes (“real” pairs) used throughout the study and were selected to maximize structural and sequence diversity while maintaining consistent length, quality, and annotation criteria (see Methods). Chain identifiers refer to the biological assembly used for model input rather than necessarily representing the first chain listed in the deposited PDB file. Several PDB entries contain multiple VHH chains binding the same antigen or multiple antigen chains interacting with distinct VHHs; in such cases, each VHH-antigen pairing is listed separately to preserve one-to-one mapping between sequences and prediction inputs. Antigen annotations follow the original PDB nomenclature and may include viral proteins, receptors, enzymes, and engineered constructs. Resolution values correspond to the experimentally determined structure used as reference for DockQ evaluation and epitope definition.

**Supplementary Table 2.** Overlapping high-confidence shuffled outliers across prediction tools. Shuffled nanobody-antigen systems (Figure 2) identified as high-confidence outliers by more than one prediction tool. Despite being mismatched (shuffled) complexes, these systems were assigned elevated ipTM scores by multiple models, although overlaps are rare and tool-dependent.

| Tool comparison | VHH PDB | Antigen PDB | ipTM (Tool 1) | ipTM (Tool 2) |
| --- | --- | --- | --- | --- |
| Chai-1 vs Boltz-2 | 4NC2 | 7R24 | 0.596 (Chai-1) | 0.813 (Boltz-2) |
|  | 7XLD | 4X7C | 0.350 (Chai-1) | 0.211 (Boltz-2) |
|  | 8EMZ | 9ETJ | 0.687 (Chai-1) | 0.578 (Boltz-2) |
|  | 9EMY | 8UKV | 0.731 (Chai-1) | 0.442 (Boltz-2) |
| Chai-1 vs AF3 | 7O06 | 5E7F | 0.572 (Chai-1) | 0.810 (AF3) |
|  | 7O06 | 7O3B | 0.705 (Chai-1) | 0.780 (AF3) |
|  | 7O3B | 7O06 | 0.716 (Chai-1) | 0.850 (AF3) |
|  | 7O3B | 8AV2 | 0.645 (Chai-1) | 0.740 (AF3) |
|  | 7TGF | 6CWG | 0.693 (Chai-1) | 0.900 (AF3) |
|  | 7TGF | 6OBO | 0.730 (Chai-1) | 0.900 (AF3) |
|  | 7TGF | 7TH2 | 0.493 (Chai-1) | 0.910 (AF3) |
|  | 7UPM | 7WKI | 0.686 (Chai-1) | 0.820 (AF3) |
|  | 7UST | 7USV | 0.688 (Chai-1) | 0.820 (AF3) |
|  | 7VNB | 8EW6 | 0.788 (Chai-1) | 0.830 (AF3) |
|  | 8FSL | 8EW6 | 0.746 (Chai-1) | 0.810 (AF3) |
|  | 8K4Q | 8EW6 | 0.785 (Chai-1) | 0.820 (AF3) |
|  | 8PIH | 8K3A | 0.381 (Chai-1) | 0.780 (AF3) |
|  | 8PYR | 6K69 | 0.653 (Chai-1) | 0.760 (AF3) |
| Boltz-2 vs AF3 | 6V7Y | 3RJQ | 0.617 (Boltz-2) | 0.620 (AF3) |

**Supplementary Figure 1.**
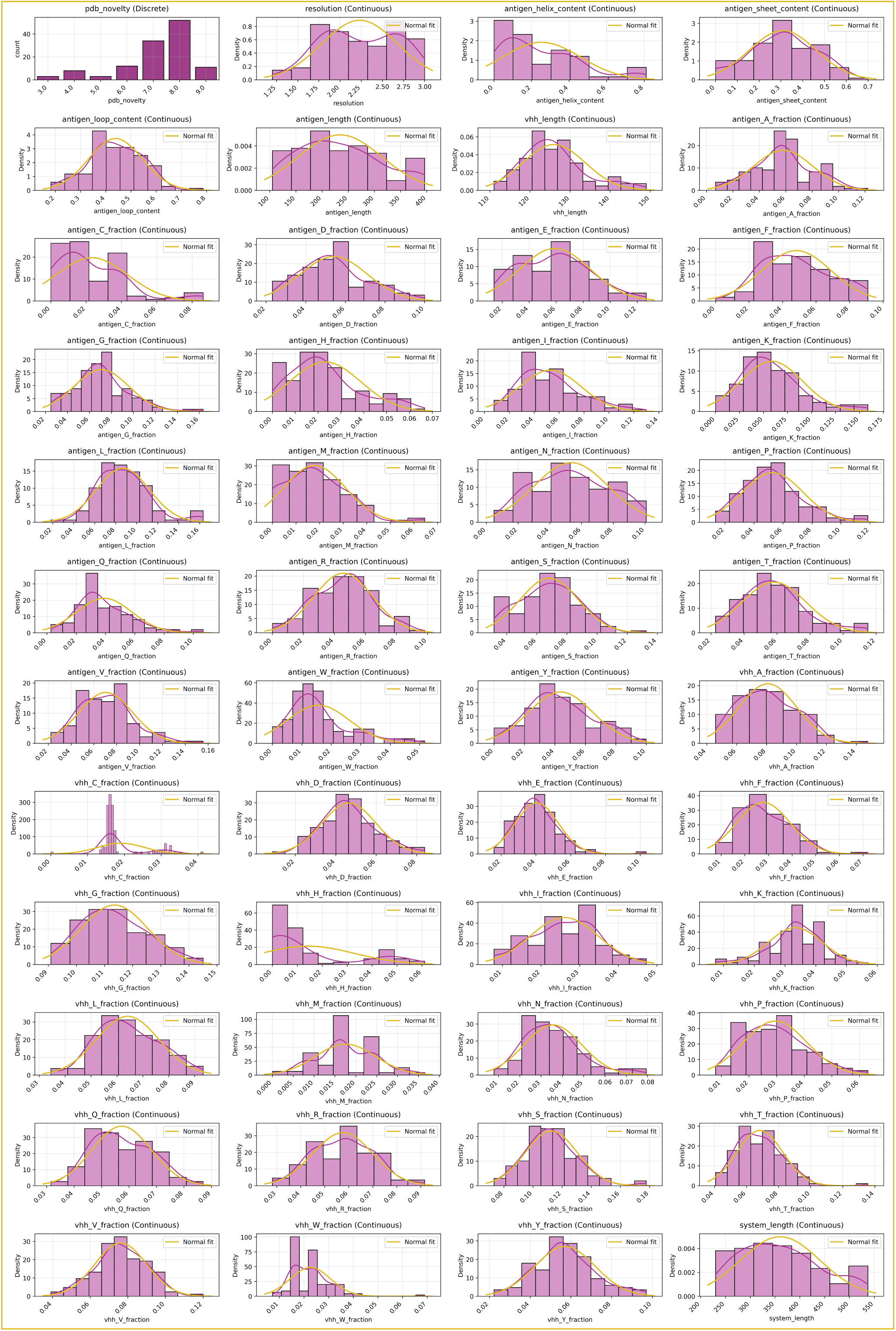
Sequence- and structure-derived feature distributions of the curated VHH-antigen benchmark dataset. Histograms summarize the distributions of structural, compositional, and sequence-derived features across all nanobody-antigen systems included in the benchmark. Each panel shows the empirical distribution (purple bars) together with a fitted normal density curve (yellow line) for visualization. Features include crystallographic metadata (PDB novelty, resolution), antigen structural composition (secondary-structure content, loop fraction, chain length), amino-acid composition fractions for antigens and VHHs, and overall system length. Antigen features are computed over the antigen chain only, whereas VHH features correspond to the nanobody sequence used as model input. Density scaling was applied to facilitate comparison across features with different numerical ranges. The broad but controlled distributions illustrate the structural and sequence diversity of the benchmark set while remaining within predefined filtering criteria (Methods), ensuring that observed modeling behavior is not driven by extreme outliers in size, composition, or secondary-structure bias.

**Supplementary Figure 2.**
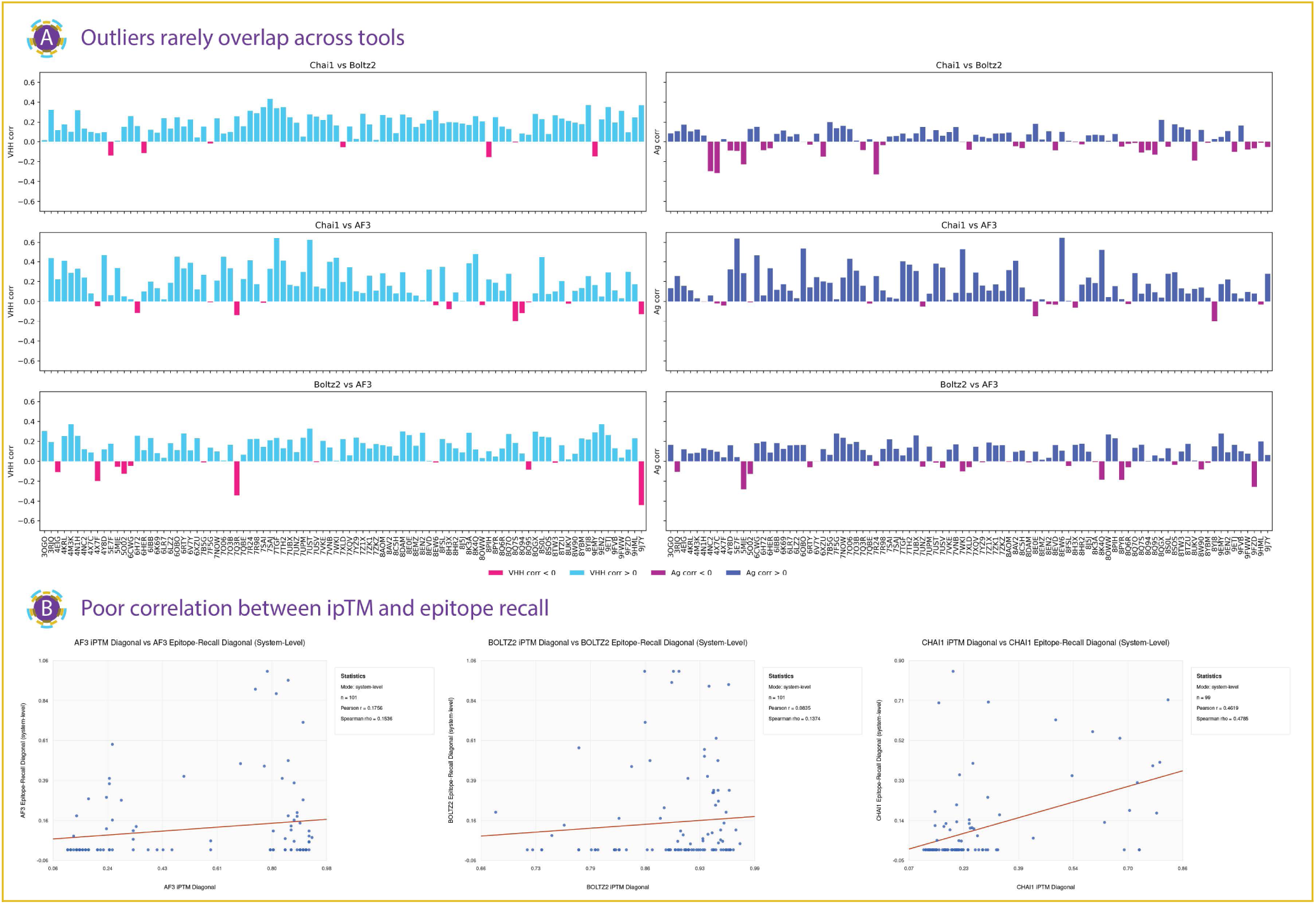
Relationship between outliers, interface confidence (ipTM) and epitope recall across prediction tools for cognate nanobody-antigen complexes.. A. *Inconsistent outliers across structure predictions tools.* Bar plots show per-system pearson correlation of predicted interaction confidence scores between pairs of tools (Chai-1 vs Boltz-2, Chai-1 vs AF3, and Boltz-2 vs AF3), computed separately for VHHs (left panels) and antigens (right panels). Positive correlations (blue) indicate agreement between tools, whereas negative correlations (pink/magenta) highlight entities for which tools assign opposing confidence trends. Across all comparisons, correlations are generally weak and frequently negative for specific VHHs or antigens, indicating substantial tool-specific disagreement. Outlier systems, where one model assigns high confidence while another assigns low confidence, are largely non-overlapping across tools, suggesting that confident failures are idiosyncratic rather than shared. B. *Poor correlation between ipTM and epitope recall.* Scatter plots show the association between diagonal interface confidence scores (ipTM) and epitope recall for AlphaFold3 (left), Boltz-2 (middle), and Chai-1 (right). Each point represents a real (cognate) VHH-antigen complex summarized as the mean value per PDB entry. The x-axis denotes the diagonal ipTM score derived from the all-vs-all interaction matrix, while the y-axis indicates epitope recall, defined as the fraction of experimentally determined antigen epitope residues recovered by model predictions according to the consensus-contact criterion described in Methods. Red lines show linear regression fits; pearson and spearman correlation coefficients are reported within each panel. AF3 and Boltz-2 exhibit weak associations between confidence and epitope recovery, whereas Chai-1 shows a stronger positive relationship, indicating partial alignment between its confidence signal and antigen contact recovery. The overall dispersion highlights that high interface confidence frequently occurs without substantial epitope recovery, reinforcing that ipTM is an imperfect proxy for specificity-relevant interface accuracy.

**Supplementary Figure 3.**
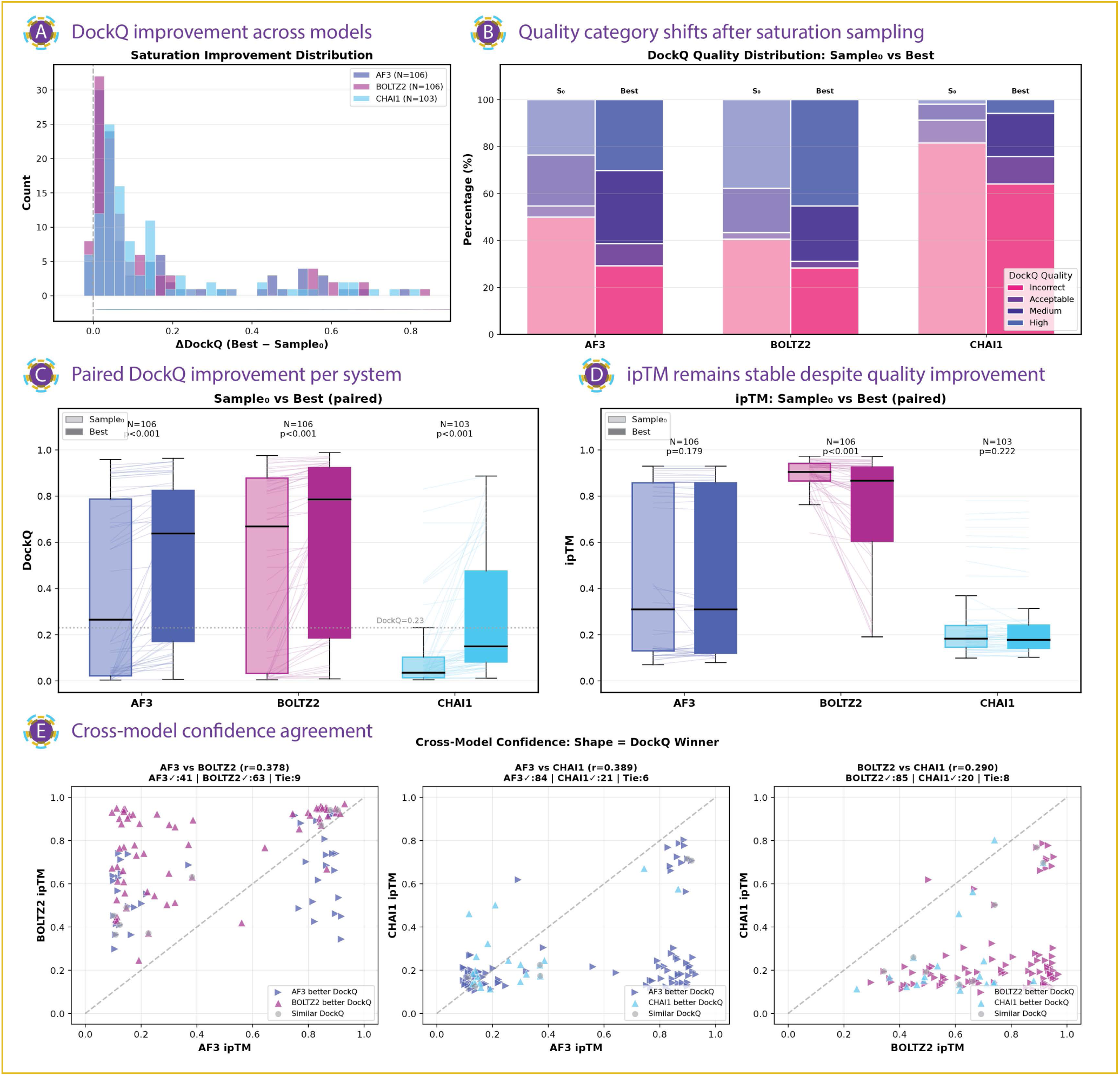
Saturation sampling improves docking quality but does not realign confidence scores across models. A. *DockQ improvement across models.* Distribution of DockQ improvement following saturation sampling, shown as ΔDockQ (Best − Sample₀) for AF3 (N = 106), Boltz-2 (N = 106), and Chai-1 (N = 103). All three models exhibit right-skewed improvement distributions, indicating that additional sampling frequently refines docking geometry. B. *Quality category shifts after saturation sampling.* DockQ quality category distributions (Incorrect, Acceptable, Medium, High) for the initial single-sample prediction (Sample₀) and the best structure obtained after saturation sampling (Best). Saturation sampling substantially increases the fraction of acceptable or higher-quality models for AF3 and Boltz-2, with more limited gains for Chai-1. C. *Paired DockQ improvement per system.* Paired comparison of DockQ scores between Sample₀ and Best predictions for each system. Lines connect paired predictions; boxplots summarize score distributions. All three models show significant improvements in DockQ after saturation sampling (Wilcoxon signed-rank test, p < 0.001). The dashed line indicates the DockQ = 0.23 acceptability threshold. D. *ipTM remains stable despite quality gains.* Paired comparison of ipTM confidence scores between Sample₀ and Best predictions. In contrast to DockQ, ipTM shows no consistent improvement with saturation sampling for AF3 or Chai-1, and only a variable decrease for Boltz-2 (p-values shown), indicating weak coupling between confidence and structural refinement. E. *Cross-model confidence agreement.* Cross-model confidence comparison. Scatter plots show pairwise ipTM values between model pairs (AF3 vs Boltz-2, AF3 vs Chai-1, Boltz-2 vs Chai-1), with point shapes indicating the model achieving higher DockQ for each system and dashed lines indicating equal confidence. pearson correlation coefficients and winner counts are reported for each comparison, revealing modest cross-model agreement and frequent mismatches between higher confidence and better structural quality.

**Supplementary Figure 4.**
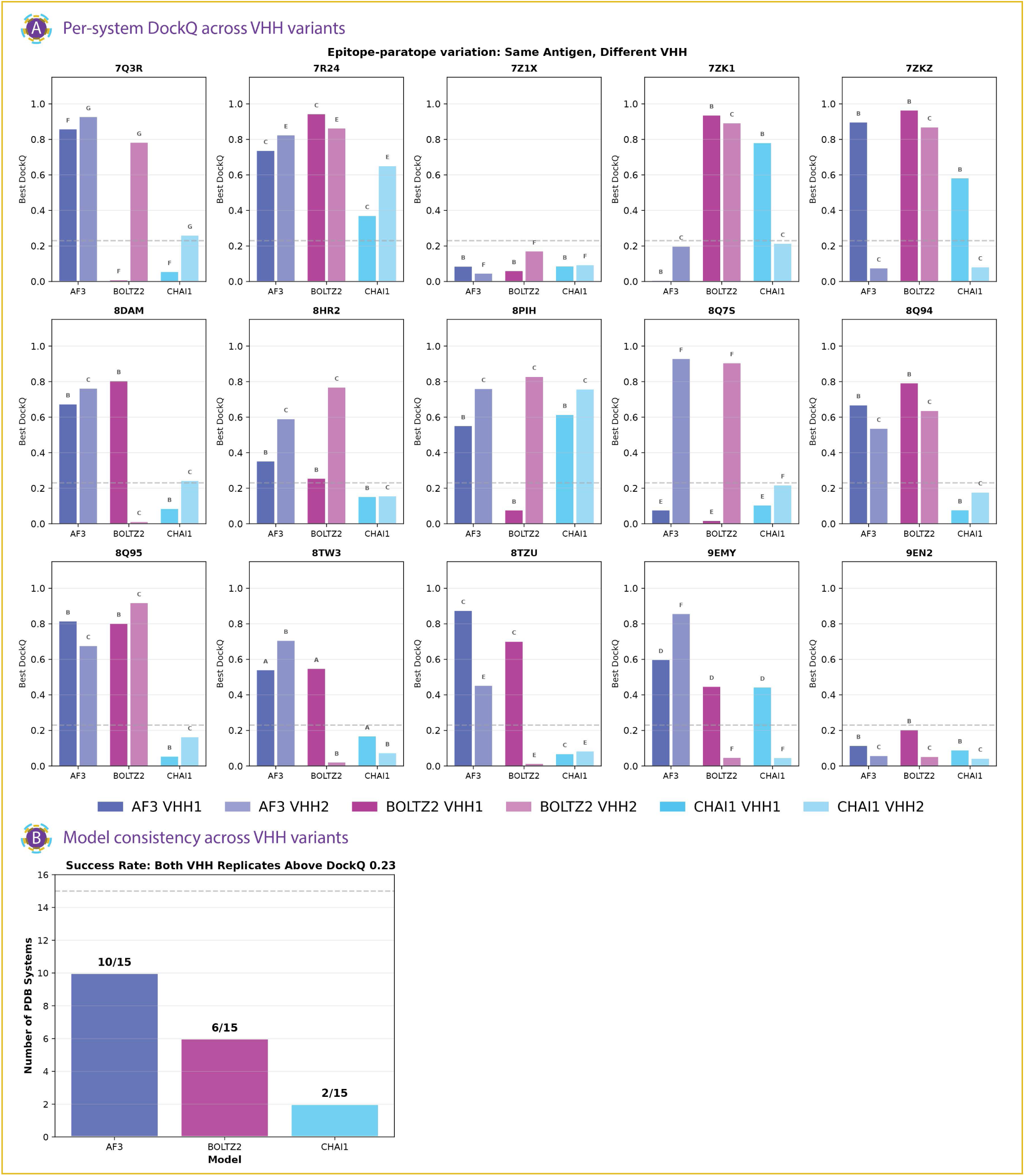
Saturation sampling improves docking quality but does not realign confidence scores across models. A. *Per-system DockQ across VHH variants.* Best DockQ scores obtained after saturation sampling for 15 antigens each evaluated with two distinct VHH binders (VHH1 and VHH2), with non-overlapping epitopes. For each antigen (PDB ID shown above each panel), bars show the best DockQ achieved by AF3, Boltz-2, and Chai-1 for each VHH replicate. The dashed horizontal line indicates the DockQ = 0.23 threshold for acceptable docking quality. Letter annotations denote relative ranking of predictions within each antigen system. High DockQ across both VHHs indicates robust recovery of the antigen epitope, whereas discordant outcomes highlight sensitivity to VHH identity. B. *Model consistency across VHH variants.* Summary of model robustness across different VHH variants. Bars show the number of antigen systems (out of 15) for which both VHH replicates achieved acceptable docking quality (DockQ ≥ 0.23). AF3 succeeds in 10/15 systems, compared to 6/15 for Boltz-2 and 2/15 for Chai-1, indicating superior consistency of AF3 in recovering correct nanobody-antigen interactions across VHH variants.

**Supplementary Figure 5.**
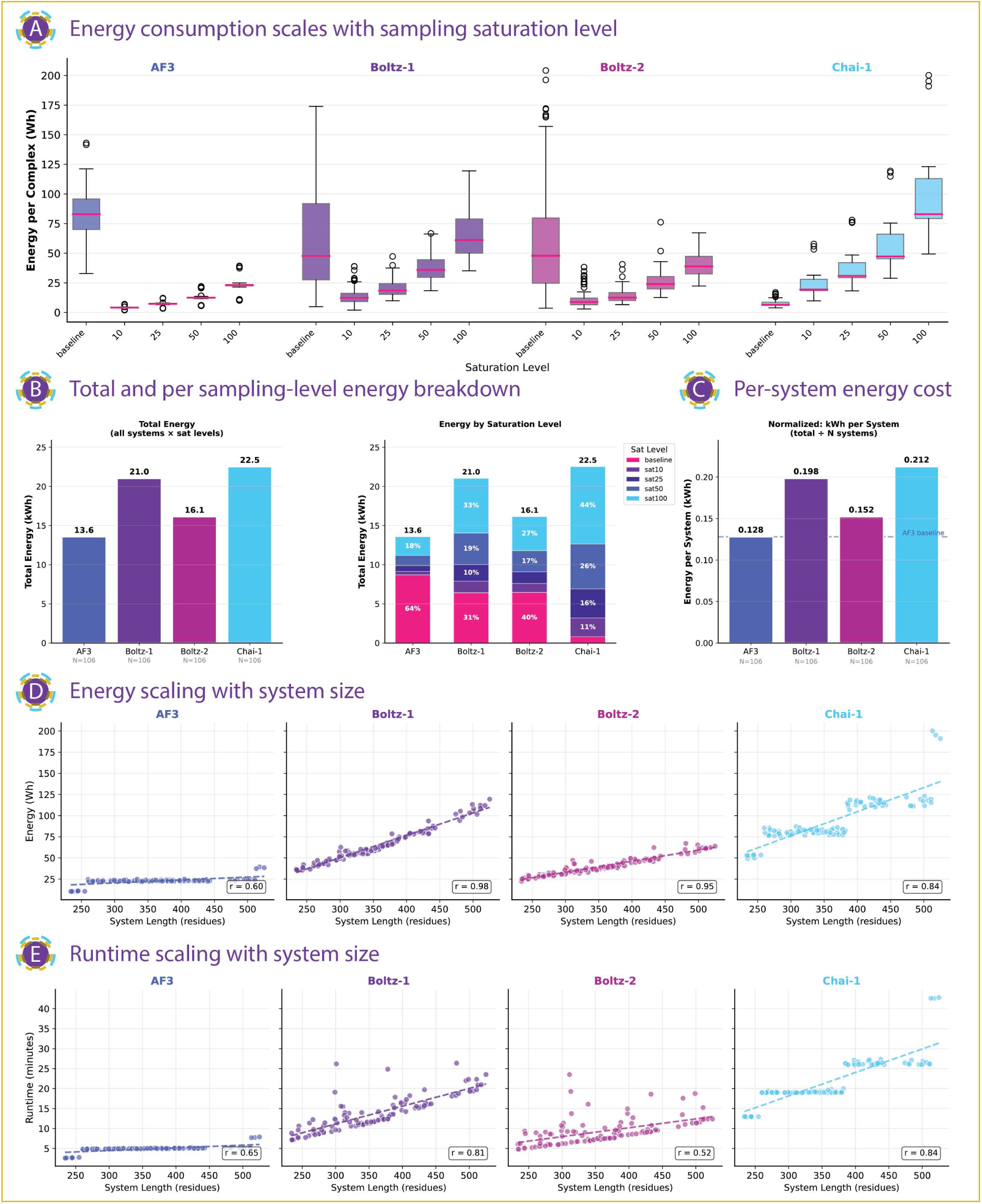
Computational cost of diffusion-based nanobody-antigen docking across models. A. Energy consumption per complex as a function of sampling depth. Boxplots show GPU energy usage (Wh) per system for AF3, Boltz-1, Boltz-2, and Chai-1 across increasing numbers of diffusion samples (baseline, 10, 25, 50, and 100). Each point corresponds to a single system; boxes summarize median and interquartile ranges. Models exhibit distinct baseline costs and marginal energy scaling with sampling depth. B. *Aggregate energy consumption across all systems and sampling levels.* Left, total GPU energy consumed (kWh) across all 106 systems and all sampling depths for each model. Middle, contribution of each sampling level to total energy usage, shown as stacked bars with percentages indicating relative contribution. C. *Normalized energy usage per system.* Bars show total energy expenditure per system (kWh) aggregated across all sampling levels, enabling direct comparison of average computational cost between models. D. *Scaling of energy consumption with system size.* Scatter plots show GPU energy usage (Wh) versus total system length (number of residues) for each model at 100 diffusion samples, with dashed lines indicating linear fits. pearson correlation coefficients (r) quantify the strength of scaling between system size and energy cost. E. *Scaling of runtime with system size.* Scatter plots show wall-clock runtime (minutes) versus system length for each model at 100 diffusion samples, with linear fits and pearson correlation coefficients shown. Runtime scaling mirrors energy scaling, with model-specific differences in slope and variance.

**Supplementary Figure 6.**
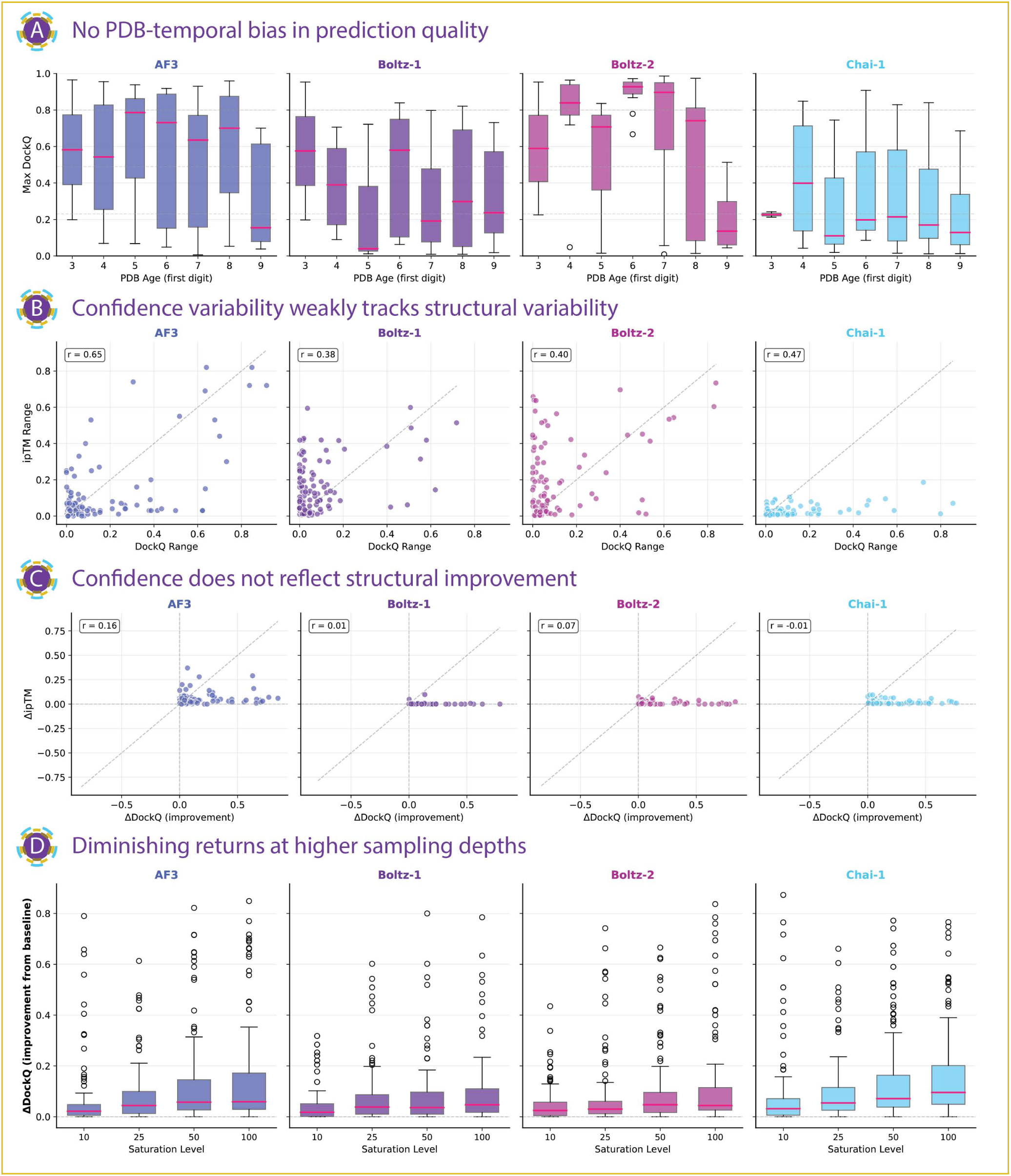
Temporal effects, confidence variability, and sampling-dependent improvements across models. A. *Prediction quality as a function of target age.* Boxplots show the distribution of maximum DockQ scores for AF3, Boltz-1, Boltz-2, and Chai-1 grouped by the first digit of the PDB deposition year. No systematic degradation in performance is observed for older structures. B. *Relationship between structural variability and confidence variability.* Scatter plots show the range of ipTM values versus DockQ scores across stochastic replicates for each system. pearson correlation coefficients (r) indicate that variability in confidence weakly to moderately correlates with variability in structural quality, depending on the model. C. *Coupling between structural improvement and confidence change.* Scatter plots show changes in ipTM (ΔipTM) versus improvements in DockQ (ΔDockQ) from baseline to the best sampled structure for each system. Near-zero correlations across all models indicate that confidence scores do not track structural refinement achieved through additional sampling. D. *Distribution of DockQ improvement as a function of sampling depth.* Boxplots show ΔDockQ relative to the baseline prediction across increasing numbers of samples (10, 25, 50, and 100). While deeper sampling increases the upper tail of achievable improvement, median gains show diminishing returns at higher sampling depths.

**Supplementary Figure 7.**
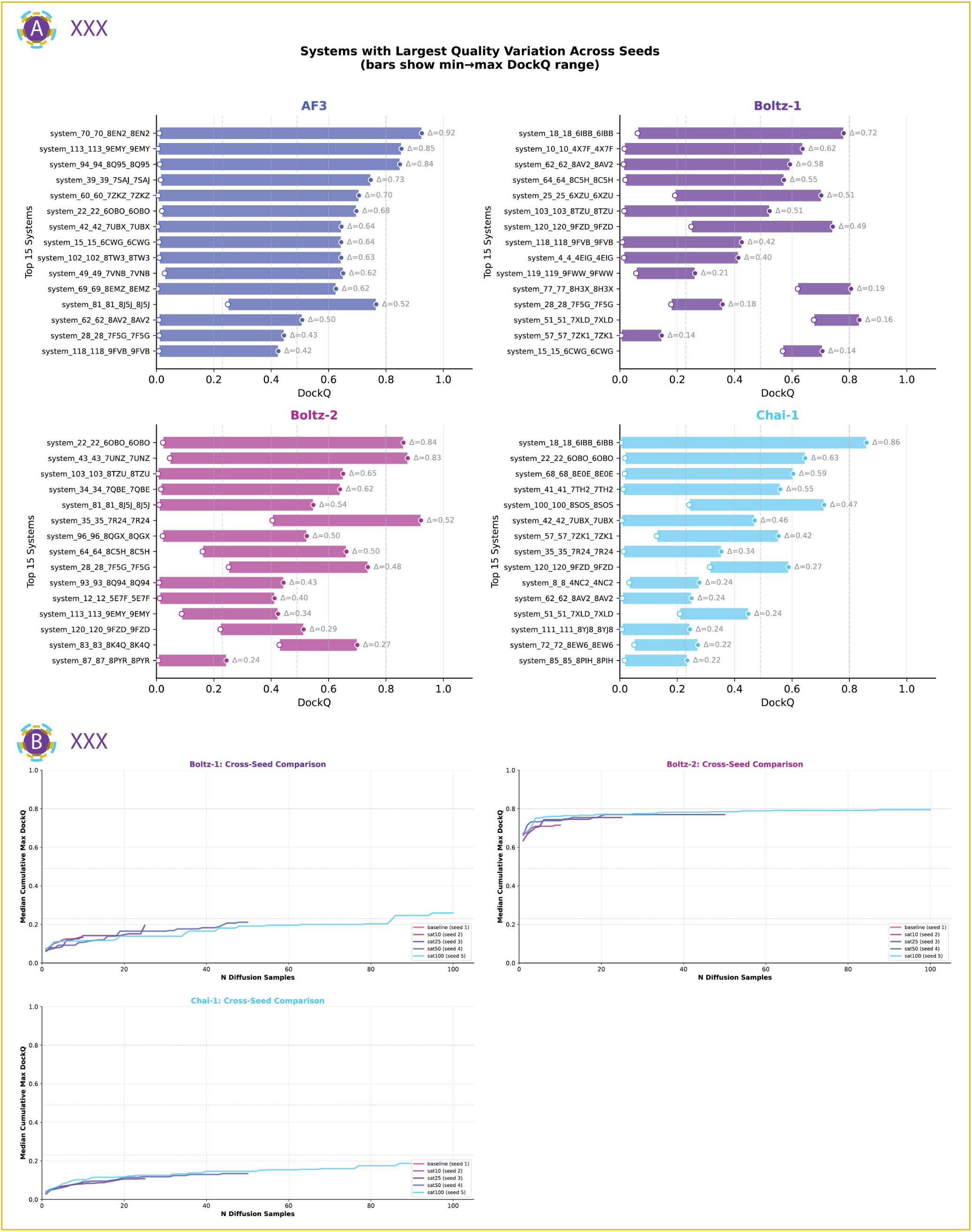
Seed-dependent variability and cross-seed convergence of docking quality. A. *Systems with the largest variability in predicted docking quality across random seeds.* Horizontal bar plots show the top 15 systems per model (AF3, Boltz-1, Boltz-2, and Chai-1) ranked by the range of DockQ scores observed across five independent seeds (Δ = max − min DockQ). Bars indicate the full min-max range for each system, highlighting substantial seed-dependent variability in achievable quality, with some systems spanning nearly the full DockQ scale. B. *Cross-seed convergence of sampling trajectories.* Line plots show the median cumulative maximum DockQ as a function of the number of diffusion samples for each seed within a model. Curves illustrate how different seeds explore distinct regions of the solution landscape and converge toward different quality plateaus as sampling increases, revealing both partial convergence and persistent seed-dependent performance differences across models.

